# Polysaccharides in cryopreservation: multidimensional systematic review of extremophilic traits and the role of selective pressure in structure-function relationships

**DOI:** 10.1101/2025.08.27.672663

**Authors:** B. M. Guerreiro, J. C. Lima, J. C. Silva, F. Freitas

## Abstract

Cryopreservation of biological matter has accumulated apex biomedical interest for its potential in elongating the shelf-life of biological matter in a state of suspended animation. In Nature, extremophilic microorganisms have an outstanding ability of surviving in habitats where extreme cold, heat, salinity and acidity defy the established boundaries for life. Through Darwinian selective adaptation, they have developed biochemical defense strategies to counteract lethal stimuli. Here, we have compiled an extremophilic EPS structure-function relationship database (XPOL-DB) which aggregates reports on 145 extremophilic and mesophilic EPS, for a total of 128 biochemical and establishes the psychrophilic chemical profile for cold adaptation. Psychrophilic EPS are highly-branched, polyanionic, elongated structures of increased flexibility and molecular weight (16–300 MDa), with predominant expression of polar monomers (GalNAc, GalA, GlcNAc, GlcA) – compared to the linear, rigid, neutral thermophilic EPS. This critical analysis revealed the significant EPS similarity between psychrophiles and halophiles suggests ice growth and extreme salinity are rooted in a shared mechanism of physical membrane destabilization, for which similar chemical traits can dualistically imbue freeze and salt tolerance. Psychrophiles are exciting testbeds for the mapping of how extreme cold selects for cryobiological EPS adaptation; and can fuel reverse engineering efforts to design optimal bio-based, non-cytotoxic cryoprotectant polysaccharides.

## Introduction

Every year, around 70–80% of Earth’s biosphere is frequently exposed to temperatures below 5 °C [1]. In Vostok, located in the East Antarctic Ice Sheet of the South Pole, temperatures can reach almost –70 °C, with maximum temperatures reaching –30 °C in the warmest months [1]. In such perilous conditions, the existence of life appears highly improbable, but nevertheless, it exists and thrives. Just as abundant life has been found in –20 °C habitats in the polar regions, Planet Earth harbors species capable of proliferating in sulfurous environments (pH 0), in soda lakes (pH 14), in the extreme salinities of the Dead Sea (6 mol/L), in high-pressure deep sea (100 kPa) and in thermal vents and hot springs with temperatures above 100 °C [2]. These species are known as extremophiles, for their ability to withstand external stressors, often considered extreme enough to defy the established boundaries for life. In cryobiology, psychrophiles are the model extremophilic strain of interest, for their ability to thrive under extreme cold. Earth’s cryosphere spans over 33 million kilometers, and although believed to be remote and inhabited, microorganisms largely populate this realm and numbers can reach those of unfrozen habitats, ranging from 10^1^–10^8^ cells/ml [1]. For such hostile environments to be so densely populated, it is largely accepted that Darwinian selective adaptation must have promoted the development of functional strategies to allow these organisms to avoid cryoinjury. One theory derived from Darwinian evolution, and coined the “Snowball Earth” hypothesis, postulates that the existence of cold-resistant microorganisms must have derived from cold-adaptive pressure rather than an intrinsic genetic disposition towards the cold. Permafrost, which constitutes about 25% of Earth’s land area and is defined as soil frozen for at least 2 years and up to 1–3 million years in the Arctic and Antarctica regions, can be incredibly biodiverse because it acts as an environmental registry of preceding time periods. The liquid medium comprising a permafrost sample encapsulates the chemical characteristics of atmospheric and geological biomes of that period, which selected for specific adaptation strategies and microorganisms [3]. With Earth having been fully frozen more than once and its latest ice age having occurred 650 million years ago [3], the geological interconnectivity between the cryosphere and other biospheres plausibly justifies why the geographical distribution of cold-adapted strains appears so ubiquitous throughout the planet. The “Snowball Earth” hypothesis further extends the same reasoning for the possibility of life in other cold planets e.g. Mars, where permafrost is equally abundant [4].

### Extremophilic species and external stressors

Microorganisms that live in extreme habitats produce extracellular polysaccharides (EPS) that enable them to withstand external stressors and survive [5]. Regardless of habitat, EPS are complex architectural macromolecules that contribute to ensuring basic functionality that a microorganism colony requires to survive: intercellular signal transduction, nutrient diffusion, molecular recognition, protection against predation, cell adhesion, biofilm formation and mechanical robustness [6]. Depending on which external stressor these microorganisms are exposed to, surviving strains have selectively adapted to possess traits correlated to the physical origin of each stimuli. Therefore, the characteristics of each EPS often expresses unique structure-function relationships relative to their extremophilic type, because their chemical nature is the product of an enabled perpetuation of life in abnormal heat, cold, salinity, acidity or pressure [6] (Table 1).

**Table 1:** Classification of the most abundant extremophilic types and their characteristics.

| Type of extremophile | Environmental stressor | Growth tolerance | Exemplary strain |
| --- | --- | --- | --- |
| Psychrophiles | Cold | < 15 °C | *Colwellia psycherythraea* |
| Halophiles | High salinity | 2–5 mol/L NaCl | *Halomonas eurihalina* |
| Thermophiles | Heat | 60–80 °C | *Thermotoga maritima* |
| Hyperthermophiles | Heat | > 80 °C | *Geobacillus tepidamans* |
| Alkaliphiles | Acidity | pH > 9 | *Acidithiobacillus thiooxidans* |
| Acidophiles | Acidity | pH < 5 | *Anaerobranca californiensis* |
| Piezophiles | Pressure | 40–60 MPa | *Shewanella benthica* |
| Radiophiles | Radiation | 5000 Gy | *Deinococcus radiodurans* |

#### Psychrophiles

Psychrophiles can grow at temperatures lower than 15 °C, some even living in sub-zero regions, such as the Arctic, Antarctic or deep-sea sediments, dominating the marine ecosystem [6]. They thrive in environments that cycle nutrients through nitrogen fixation, nitrification and denitrification, photosynthesis, sulphur redox reactions, methanogenesis and organic compound transformation [7], and can be found in snow, glaciers, floating ice sheets and permafrost [3]. The common factor between all these geological structures is their submersed hydrological connectivity, which serves as probable basis for bacterial migrations throughout various ecosystems that appear physically isolated. Unlike other extremophilic types, psychrophiles exhibit increased EPS production at low temperatures, despite a general reduction in metabolic activity. This suggests EPS may serve as a reservoir for nutrient accumulation or play an active role in cold adaptation [8–10]. Psychrophilic EPSs have shown embedded ice-active proteins in their structure to increase their affinity for ice binding, and the psychrophilic host cell wall can function as an ice-nucleating particle, contributing to snow formation [11].

#### Other extremophilic types

Halophiles can withstand hypersaline environments containing 5–20% w/v salinity content [6], because their halophilic EPSs often contain elevated amounts of uronic acids and sulphate groups. These anionic moieties allow for great heavy metal binding and gelling capacity, the latter allowing to regulate diffusion and osmotic pressure. Thermophiles are able to live at temperatures above 55 °C and inhabit marine and terrestrial hot springs, and deep sea thermal vents [6]. They produce thermostable polymers, able to withstand high degradation temperatures up to 280 °C [12]. Acidophiles have optimal growth at pH < 3, cannot withstand neutral pH environments [6]. Sulfidic mine areas, marine volcanic vents and pyrite rocks are optimal for their bioleaching activity due to the beneficial presence of sulfur, sulfide and their oxidates, and valuable metals like Fe, Cu, Co, Al, Mg, Zn and Mn. Alkaliphiles are adapted to environments of pH > 9, often between 10–12 [6], such as soda lakes, deserts or historic limestone. Some alkaliphilic EPSs actively dissolve calcite during biofilm formation and accumulate unusual carbon sources like acetate [13].

### Biological limits to life

This overarching endurance to the most extreme habitats has challenged previous tenets of biological limits, but in cold-adapted psychrophiles two unavoidable constraints have been identified: water availability and the glass transition temperature (T_*g*_). Although water in liquid form at –56 °C has been found in glacial ice as thin films or brines [14], life requires bioavailable water to exist and participate in metabolic reactions [15]. Water activity (*a*_*w*_) is a fractional measure of bioavailable water and the minimum threshold for the occurrence of life processes has been identified at *a*_*w*_ = 0.61 [15]. In the salt brines of McMurdo Dry Valleys in Antarctica, no microbial activity has been found although water films are present all year (*a*_*w*_ = 0.45). However, bioavailable water also implies an increased ice growth probability. In Nature, geographic cooling rates are very slow and intracellular ice formation (IIF) rarely occurs because the cytoplasm becomes densely packed due to an osmotic efflux of water to the external environment. Thus, extensive extracellular ice formation (EIF) is dominant. This adaptation by dehydration leads to internal vitrification and implies that the source of damage is usually dehydration pressure [16]. Therefore, the ability of a biological system to vitrify into an amorphous glassy state (at a given T_*g*_) so viscous that molecular reorientation hinders ice formation, slows down metabolism to geological time scales [17] and leaves the organism in a state of suspended animation, has been considered the second limit to life. The lowest natural T_*g*_ found was that of *Arthrobacter arilaitensis* at –26 °C [18], while the lowest artificial T_*g*_ found was that of *Lactobacillus delbrueckii* sp. in the presence of 0.58 M dimethyl sulfoxide (DMSO) at –51 °C [19]. Microbial activity in permafrost ranging between –17 °C [20] and –25 °C [21] falls under these constraints.

### The Cryosphere

The cryosphere, the domain of Earth’s surface which concerns or connects to geographical regions where water is in solid form, is mostly populated by Proteobacteria, mostly predominant in cold seawater and glaciers, but found ubiquitously in both Earth poles and all cold climates in between. Actinobacteria and Firmicutes are found in permafrost due to their dormant-inducing [3] and spore-forming [22] properties (Table 2). *α*/*γ*-proteobacteria dominate all regions of the Artic connected to a liquid geography or where water percolation exists, namely seawater, sea ice and ice wedges. *β*-proteobacteria prefer to inhabit Antarctic glacial ice, possibly because the longer summer seasons in Antarctica and drier conditions also lead to reduced freeze-thaw cycles, preserving the solid habitats for longer [3]. Heterotrophic *γ*-proteobacteria (g. *Glaciecola, Colwellia*) and Flavobacteriia (genera *Polaribacter, Psychrobacter, Psychroflexus, Flavobacterium*) prefer sea ice [1]. Sea ice accumulates denser quantities of dissolved organic matter and supports heterotrophy. Cryoconite holes are dominated by Proteobacteria and Actinobacteria, but also contain members of the genera *Pseudomonas, Polarimonas, Micrococcus, Cryobacterium* and *Flavobacterium* [23]. The *γ*-proteobacterium *Colwellia psychrerythraea* 34H is of particular interest to cryobiology. It lives in sea ice brines, is found in both the Arctic and Antarctic poles [24], grows at extreme conditions of –12 °C and 16% salinity, is motile in viscous media and over-produces EPS when enclosed into the ice matrix [9]. It secretes a cold-active aminopeptidase, responsible for nutrient capture, that is further stabilized by EPS presence [25]. Its capsular polysaccharides (CPS) contain regularly spaced threonine residues that structurally mimic ice binding proteins (IBP), therefore facilitating the colonization of sea ice [26, 27]. With regards to the “Snowball Earth” hypothesis, the strain 34H has shown to encode genes from ice-active bacteriophages and distant organisms [28], demonstrating that cold adaptation due to mid-Eocenic extensive ice formation must also derive from horizontal gene transfer [29].

**Table 2:** Psychrophilic incidence profile based on cryosphere region.

| Cryosphere biome | Dominant phylum |
| --- | --- |
| Seawater | α-proteobacteria |
| Snow, glacial ice, and subglacial Lake Whillans | β-proteobacteria |
| Sea ice | γ-proteobacteria and Flavobacteria |
| Ice wedges in permafrost | γ-proteobacteria |
| Permafrost | Actinobacteria, Firmicutes and Δ-proteobacteria |

### A primer on ice physics and cryobiology

To contextualize the role of selective pressure on the structural adaptation of psychrophiles to cold stimuli, it is beneficial to briefly review the microscopic mechanisms of nucleation and ice growth modulation that regulate how a macroscopical expression of increased post-thaw survival may emerge. This understanding allows to reason why certain phenotypical adaptations in psychrophiles naturally emerge.

#### Fundamentals of ice growth

The process of freezing, in which ice crystals form, involves water molecule aggregation into periodic, organized clusters: crystals. In pure water, water molecules co-interact by means of their negatively charged oxygen with the hydrogen of a neighboring water molecule: H–O–**H** *…***O**–H. Effectively, these are hydrogen bond donor-acceptor dynamics, so any molecule capable of providing the same electrostatic interaction with the hydrogen atoms of water is effectively a disruptor of this periodic structure. Therefore, the existence of a permanent dipole moment is one of the most crucial features of cryoprotective agents (CPAs). The gold standard example is DMSO, where the negative charge delocalization towards the oxygen in the S=O bond establishes a similar dipole moment to that of the oxygen in the O–H bonds of water [30]. The result is a disorganized ice crystal structure, with DMSO acting as molecular inclusions that fragilizes the crystalline structure.

#### Small-molecule & polymeric cryoprotectants

Small-molecule cryoprotective agents (CPAs) like synthetic permeable glycerol and DMSO [31] or naturally excreted impermeable sugars like trehalose [32, 33] can efficiently mitigate ice growth or exert IRI [34]. From alcohols, sugars, sulfoxides, amides and amines, currently a total of 20 small molecule CPAs are consistently considered to have some magnitude of a cryoprotective effect on a large variety of biological settings [35]. A thorough description of several cryoprotectant agents and their clinical applications can be found elsewhere [36]. However, several polyols [34, 37], polyampholytes [38–40], and antifreeze proteins [41–44] have demonstrated enhanced benefits over small CPAs. At the polymer scale, high molecular weight, complex monomer composition, spatial conformations, stereoregularity, polyionicity and increased viscosity can have a compound effect on water diffusion, fluid rheology, ice thermodynamics and bulk/interfacial water physics, properties which small CPAs do not present unless very high and toxic concentrations are used [45]. For example, an induced viscosity increase by polymeric CPAs reduces osmotic shock by loss of ionic diffusivity, and the chemical nature of their charged side chains has been reported to be the primary indicator of cryoprotective potential [46]. Polyampholytes with a proper balance of hydrophilic and hydrophobic moieties have shown to enhance the survival rate of cryopreserved biologicals [47]. If mostly hydrophobic, polyampholytes contribute to lowering the T_*g*_ of the bilipid membrane [48], but post-thaw recovery of A549 cells and red blood cells was only enhanced if mostly hydrophilic [49]. The hydrophilic-hydrophobic structural balance has been validated for antifreeze proteins and polyampholytes and shown to be a major driver of thermal hysteresis and dynamic ice shaping [50], but increased hydrophobicity has also been correlated to greater IRI and increased affinity to cell membrane interactions that further allow volumetric regulation, thus averting cell death [48]. The role of a hydrophobic character in cell membrane interactions has also been extensively noted. Amphiphilic [51] and cationic [52] polymers can increase membrane permeability, alginate-RGDS peptides can improve permeability, cell recognition and mitigate apoptosis [53], and dextran-based polyampholytes with increased amination lead to greater concomitant ice binding and cell surface adsorption [54]. Trehalose-decorated polymers suggested that the epitaxial location of the cryoprotective moiety is crucial from a macromolecular perspective. If trehalose is embedded in the side chain, it can stabilize proteins under heat and freeze-dry stress, but their action in cells is unknown [32, 55]. If trehalose is part of the backbone, some recovery post-thaw was observed, but resulted in lower membrane integrity than with standalone DMSO [56]. Other sparse accounts [6] of polyxylomannans being able to modulate ice growth [57] and fucose-rich polysaccharides [58–63] showing promising cryoprotective evidence has led to the proposition that naturally secreted EPS have an inherent ability to protect their hosts against cold stressors.

#### Mechanisms of action

*Cryoprotection* has often served as an umbrella term for diverse mechanisms of cryoprotective action convoluted solely under a biological post-thaw benefit. A substance capable of preserving biological matter at sub-zero temperatures is said to be cryoprotective, but the physical mechanisms by which biological success arises may vary extensively. This terminology is often ingrained in the literature and has been generalized, but in fact, cryoprotective function may be claimed by physical observation of ice modulation, in non-biological settings. Several mechanistic variations of ice growth disruption have been reported: direct ice growth inhibition [64], supercooling [30], ice recrystallization inhibition (IRI) [65], nucleation inhibition [46, 66], nucleation promotion [67], thermal hysteresis (TH) [68], among others. In general, the ability of a molecule to interact strongly with water often means it can efficiently disrupt the molecular directionality and crystal periodicity required in nucleation and ice growth. From a mechanistic standpoint, ice growth hindrance occurs at the quasi-liquid layer (QLL), which delimits the interface between a growing solid ice front and the unfrozen water fraction [69] (Figure 1). Given the heat and mass transfer properties of a growing front, thermodynamic processes like dynamic ice shaping, growth inhibition and any sort of chemical potential changes from surface energy modifications can only efficiently occur on a phase where molecules can diffuse, i.e. where kinetic energy is predominant, hence why most of these phenomena occur at the QLL interface and not in the ice bulk [70].

**Figure 1:**
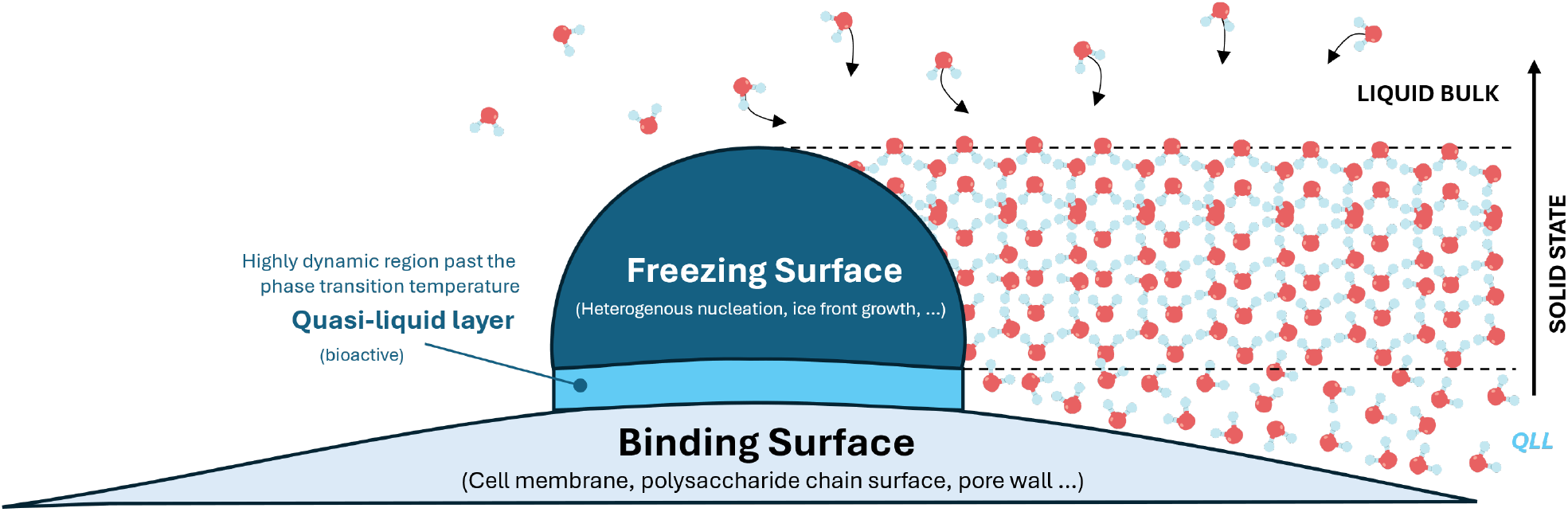
The QLL interface is a thermodynamically active region of transient water-ice phase transformations, highly relevant to cryoprotective mechanisms of action.

#### The adsorption-inhibition model

The core central description of how macromolecules may influence ice growth is largely inspired by the adsorption-inhibition mechanism of antifreeze proteins (AFPs), the first model accounting for ice growth disruption and cold-adapted survival [71]. During the late 1960s, it was initially believed that the ability of Antarctic fishes to survive and tolerate supercooling temperatures from 6 °C down to –2.5 °C [72] arose from freezing point depression of blood serum by sodium chloride, urea or free amino acids [73]. Later, it was found that it was because of AFPs present in the blood that could adsorb to the planar facets of crystals and inhibit ice growth [74]. AFPs can be structurally very diverse, but the simplest example is the type I AFP from the winter flounder *Pleuronectes americanus*. This AFP is a simple 37-residue helix [75] composed of one negatively charged threonine residue separated by seven neutral alanine residues. The threonines, aptly spaced to mimic the 16.7 Å separation of hydrogen bonds in ice [76], actively bind to the non-basal plane (*a*-axis) of the ice crystal lattice by reversible Langmuir adsorption [77], which generates thermal hysteresis and dynamically reshapes the ice crystals from a native hexagonal shape to bipyramidal crystals [78]. This modulation of ice curvature is known as the Kelvin, or Gibbs-Thomson ice binding effect [77]. At the QLL between the AFP and the ice front, further access of new water molecules to the ice front is precluded, because AFP adsorption generates steric hindrance, and an increased interfacial tension of the sharper bipyramidal vertices reduces chemical potential for ice growth (Figure 2). The adsorption-inhibition model is currently the best proposal of how AFPs induce the supercooling of water from protein-ice interactions, and has inspired the search of adsorptive inhibitory behavior in polysaccharides as a strong potential indicator of cryoprotective activity. So far, similar trends have been observed for polyampholytes [38–40] and polysaccharides [61, 63].

**Figure 2:**
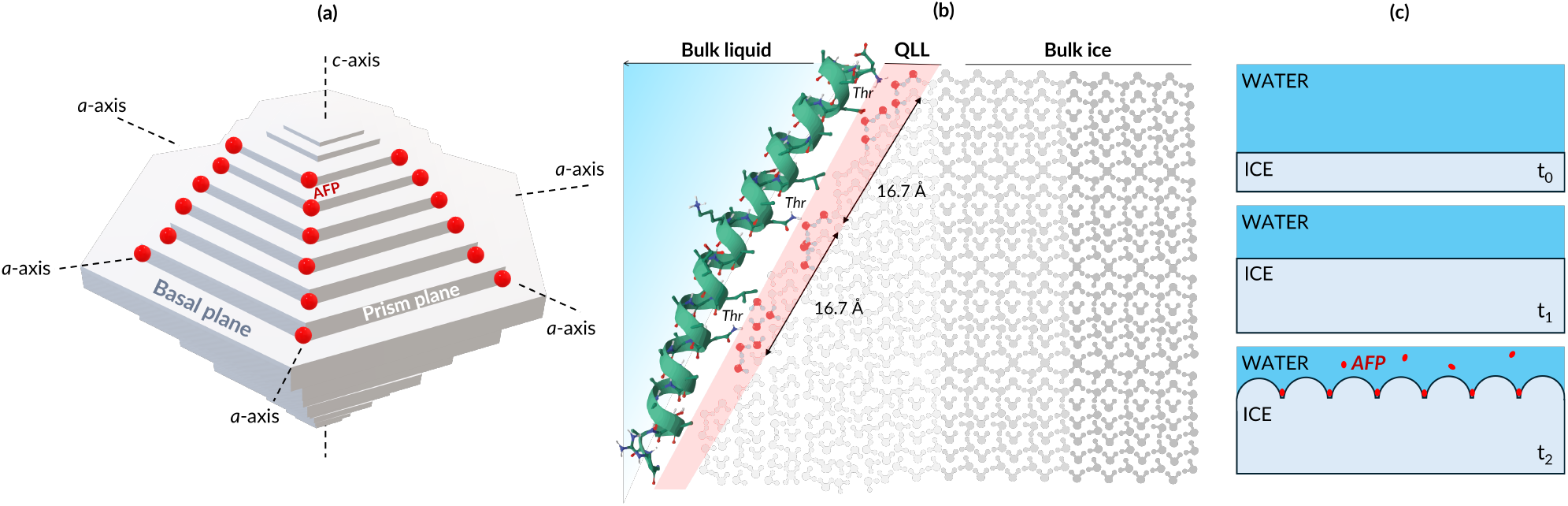
The adsorption-inhibition mechanism represented at different length scales: (a) ice crystal, (b) adsorption interface and (c) macroscopic ice front. At subzero temperatures, hexagonal ice crystals undergo dynamic ice shaping to bipyramidal crystals as AFPs bind to the a-axis of the ice lattice (a). A close-up to the ice lattice reveals that active inhibition occurs at the QLL (red band), when the AFP (green ribbon) adsorbs to the growing ice front (gray). Bioactive threonines (Thr, blue) bind to QLL water molecules (red) and exert ice front curvature and reshaping, increasing surface tension and precluding the migration of bulk water to the ice front (b). At the macroscale, this phenomenon can also be observed in 2D directional freezing, whereas the stepwise advancement of the ice front is inhibited by AFP binding to specific kink sites (red), forming dendritic ice as time elapses (*t*_0_ *→ t*_1_ *→ t*_2_) (c).

### Biological evidence of cold adaptation

Psychrophiles exhibit multifaceted adaptations to cold environments, where low temperatures reduce molecular motion, water availability, and membrane fluidity; and drastically increase intercellular viscosity by solute supersaturation and mechanical stress to the cell boundary during osmotic shock [79].

#### Cellular substructures, DNA, lipids, proteins, enzymes

Briefly, psychrophiles have adapted to (i) embed flexible polyunsaturated fatty acids (PUFAs) of higher melting point into the cell surface (e.g. eicos-apentaenoic acid) to be able to undergo membranar liquid-to-gel phase changes and increase fluidity, (ii) implement more active transport membrane proteins to maintain hypothermic homeostasis e.g. regulators of glycine-betaine turnover for protection against hyperosmotic shock [80–82]; (iii) ensure DNA replication, transcription and protein synthesis using dedicated cold-shock proteins that chaperone the hypothermal process [83–85]; (iv) produce intracellular and extracellular enzymes with flexible tertiary structures capable of hypothermic catalysis [86] due to their higher flexibility, disorder, dominant hydrophobicity [87], and reduced activation energies [88] and thermal stability [89] and (v) synthesize AFPs, IBPs, cold-shock proteins, and other cryoactive glycoproteins. Detailed reviews on each of these can be found elsewhere, whereas this review will focus on the production and accumulation of EPS, which can act as cryoprotectants and osmoprotectants [90].

#### Polysaccharides

A psychrophilic EPS has peculiar features that contribute to cryoprotection [6]. The most basic evidence of it is the psychrophilic *γ*-proteobacterium *Colwellia psychrerythraea* 34H demonstrating better survival rates after –80 °C deep-freeze when exposed to its own EPS rather than common cryoprotectants like glycerol [9]. Psychrophilic EPSs have a monomer composition often rich in carboxyl and phosphate groups which confer negative charge, and other uronic acids that are overexpressed at lower temperatures. Polyanionicity favors ice binding but also heavy metal binding; and a high molecular weight (M_w_) contributes to increased water binding capacity and favors tertiary structure formation (greater entanglement and increased binding sites) that enhances the cryoprotective effect. In terms of structure, helicoidal arrangements have been shown to promote IRI [91], and a structural repeating unit (SRU) composed of a balance of hydrophilic and hydrophobic monomers that corresponds to crystal spacings was shown to be optimal [91], rather than fully hydrophilic structures. Besides a cryoprotective effect, some psychrophilic EPSs provide a buffering effect against high salinity (osmoprotection) as well [92], as certain geofeatures found in ice structures such as frost flowers, cryoconite holes and under-ice algal mats contain a combination of both stressors [93]. These EPSs can influence the physics of the ice matrix, aid in external attachment, and promote cell aggregation and biofilm formation both in and under ice mats [1]. Thermostable psychrophilic EPSs can also protect proteases from thermal denaturation [6]. This ecological evidence is in alignment with the extensive documentation of osmoregulatory [94, 95], anti-inflammatory [94, 96, 97], antitumoral [98, 99], immunoregulatory [97, 100], wound healing [101,102], gel-forming [96, 103], emulsifying [104–106] and neuroprotective [107] effects shown *in vitro*, among others, that further support the broad use of EPS in the cryopreservation of complex biological systems, as they can synergistically protect biological systems on different fronts.

### Main Contributions of this Review

EPS produced by extremophilic microorganisms have been considered exciting testbeds for large-scale bioreactor production and clinical application [6] but their characterization remains challenging due to very complex structures [91] and poorly understood structure-function relationships. With over 100 possible monomeric sugars that may constitute a polysaccharide, the possible combinations of composition, molecular weight, secondary and tertiary structure, global charge, binding regions, stereoregularity and chemical affinity that play a concerted role in function largely outweigh those in proteins. The glycosidic linkage also has less angular constraints than the peptide linkage as it lacks a double bond character, which magnifies conformational flexibility. However, it has been acknowledged in the last decade that the information regarding the structure-function relationships of EPS, mainly those psychrophilic, must be further deepened to reveal optimally advantageous structures for protection and stabilization effects [6]. Therefore, the goal of this review is twofold. First, we embarked on systematically examining how environmental stressors shape the compositional and structural features of extremophilic EPS, based on the available extremophilic polysaccharide literature, which we curated into a structure-function relationship database (`XPOL-DB`). We posited as ground-truth hypothesis that a biological adaptation response to different stressors would result in distinct phenotypic expression to ensure survival. For psychrophiles, freezing damage would exert selective pressure that promotes the overexpression of certain advantageous EPS attributes over others. Second, we critically approached the available data in order to generate additional hidden insight by generating compositional fingerprints for different types of extremophiles. By comparing and visualizing spectrally opposite stressors in terms of which exclusive polysaccharide traits emerged in each (e.g. thermophiles vs. psychrophiles) and contrasting those with non-extremophilic strains (mesophiles), we differentially pinpointed which compositional and structural attributes describe the cryoprotective effect. The extensive mapping and understanding of psychrophilic polysaccharides provides a bio-inspired approach towards advancing the emerging field of polysaccharide-based cryoprotection.

## Materials & Methods

### Extremophilic EPS: literature data collection

A relational database of extremophilic polysaccharide research, `XPOL-DB`, was compiled from an extensive data scraping of all literature containing a chemical characterization of extremophilic EPS that possessed a biological function of relevance. In total, 145 extremophilic and mesophilic EPS were reported, for a total of 144 attributes (128 organically reported and 16 mathematically calculated parameters). The database was split into nine different categories clustering several attributes: host identity (12 attributes), EPS growth (22), monomeric composition (33), structure (12), macromolecular fractions (10), physicochemical properties (15), biological function (18), biological evidence of cryoprotection (7) and evidenced mechanisms of action (7). The full database, partially described in Table *SI*.*1*, and supporting documentation necessary for the multidimensional analysis performed herein can be found [online here](https://docs.google.com/spreadsheets/d/1Pau7IhcXFqHlOs0PBtbPr0CBMyV3wPbn/edit?usp=sharing&ouid=110571406079464345723&rtpof=true&sd=true). All data computations, calculations and plotting were performed in Jupyter Notebook v7.3.2 using open-source Python v3.7 data visualization library dependencies.

### One-hot encoding

A categorical parameter *p* with *m* different possible values can be numerically encoded using binary values, generating *p*_*m*_ different attributes one-hot encoded as 1 or 0. A one-hot encoding algorithm was performed on categorical parameters reported for biological function and glycosidic linkage types, in order to generate numerically computable data for the correlation matrix plots produced in Figures 12 and 15.

### Statistical analysis

All statistical analysis was performed in GraphPad Prism 8.0.1. Results are shown as mean ± standard deviation (*σ*^2^). Statistical significance between two sampling populations was assessed with a t-student statistical test, or a one-way ANOVA with Sidak’s multiple comparisons for multiple population comparison. Whenever applicable, the *p*-value significance threshold, determined for a 95% confidence interval (CI, *α* = 0.05), was reported using The New England Journal of Medicine (NEJM) symbology: 0.12 (ns, not significant), 0.033 (*), 0.002 (**), *<*0.001 (***).

## Results

### XPOL-DB: a structure-function polysaccharide cryopreservation dataset

A structure-function relational polysaccharide database was compiled, aggregating a total of 128 organically reported parameters in known extremophile literature, for a total of 145 EPS structures from bacterial, algal and fungal species. Each polysaccharide contains a thorough description of its producing microorganism characteristics, the optimal biometrics conditions for microorganism growth and EPS production, the polysaccharide monomer composition and structure, the macromolecular fractions (i.e. protein content, lipid content) of the secreted EPS, its physicochemical properties and the biological functionalities of each polysaccharide, in particular regarding cryoprotective evidence and potential mechanisms of action. The multidimensional analysis of the `XPOL-DB` database was rationalized as a binary classification problem, wherein all collected parameters acted as predictor variables to infer a cryoprotective outcome, to tackle the primary research question:

> *“Which parametric trends contribute to a polysaccharide expressing cryoprotective action?”*

By grouping each polysaccharide into an extremophilic class and creating extremophilic profiles based on unique characteristics, structural trends and monomer composition fingerprints of their representative EPS, one can derive insights of high statistical significance when contrasting e.g. psychrophiles, thermophiles or halophiles. In other words, the features observed in cold-adapted psychrophiles and unobserved in thermophiles, may serve as conduit for establishing which features are contributive for the expression of cryoprotective activity. Figure 3 describes the data eligibility rationale and reports the final data inclusion based on these constraints, as well a surface-level overview of the database meta-data. Given the strong correlation between the chemical properties of a molecule and its cryoprotective potential, the only foundational requirement in this database that prompted an exclusion process was that the monomer composition of a given polysaccharide must be reported, otherwise no functional insights could be derived from relational data. The `XPOL-DB` is comprised of seven extremophilic classes (Figure 3b). Psychrophiles constitute the majority of polysaccharide entries (41.4%) due to their study relevance but were evenly balanced by 22.1% halophiles and 16.6% thermophiles, to avoid inferential bias. The remaining classes accounted for the remaining 19.9%: halothermophiles (4.2%), haloalkaliphiles (2.1%) and alkaliphiles (2.8%); and mesophiles (11%), a non-extremophilic class. The significantly lower representation (one-fifth of the dataset) of most extremophiles compared to psychrophiles, and thermophiles is due to scarce research and not biased heterogeneity. The specific external stressor for each extremophilic class has been described in Table 1. Figure 4 summarizes the phylum, habitat and geodemographic distributions of extremophilic polysaccharides according to their class. Proteobacteria (53.9%) and Firmicutes (32.4%) are the dominant phyla. Not only do they produce almost 90% of all polysaccharides reported, but they ubiquitously represent all extremophilic types and show extensive adaptability to a wide range of habitats. Psychrophiles and halophiles are mostly Proteobacteria, while thermophiles are predominantly Firmicutes.

**Figure 3:**
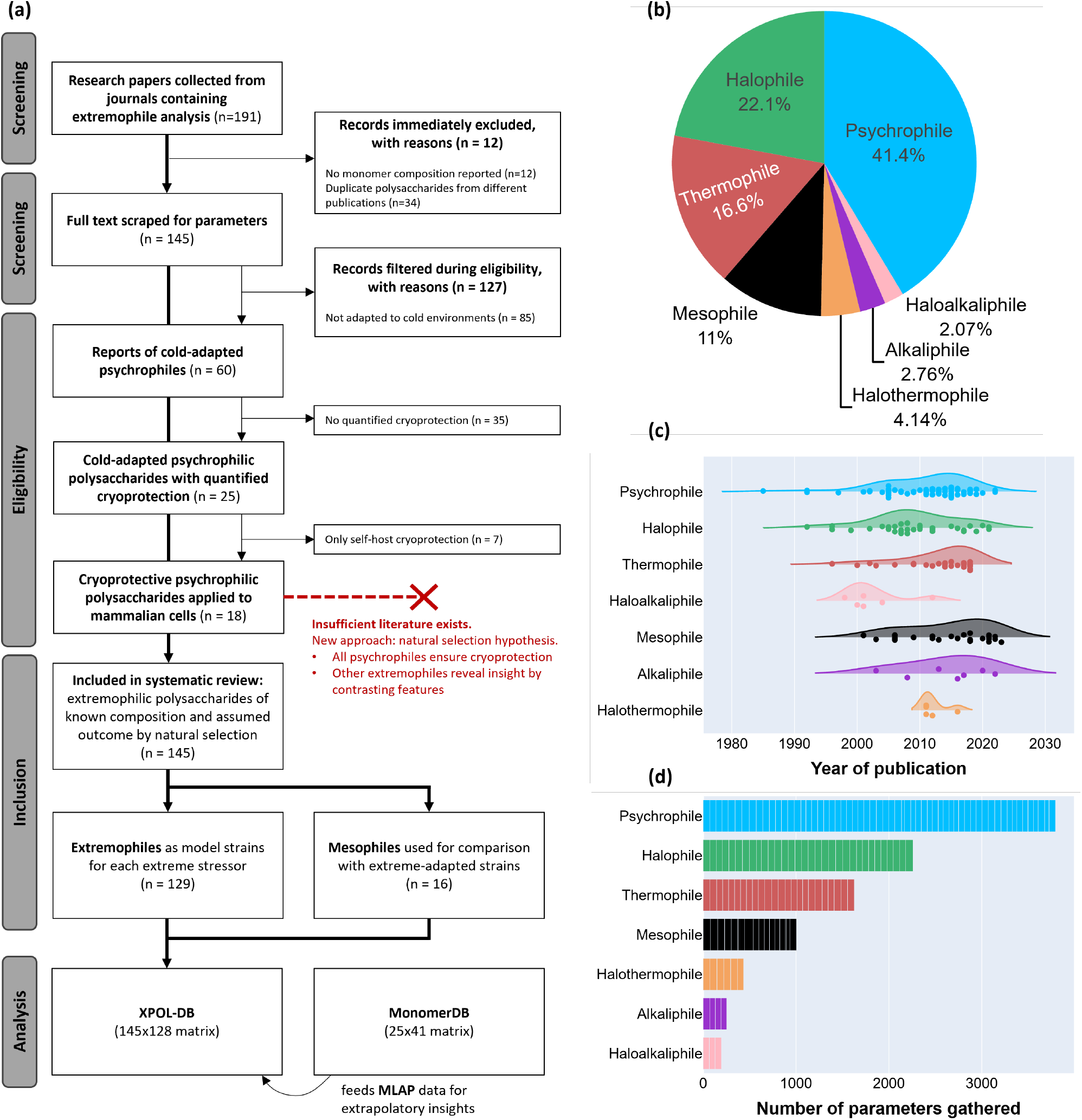
Metadata of the extremophilic polysaccharide database. (a) Decision tree for data collection, screening, eligibility and final data inclusion. The bold arrow represents the adopted pathway from screening to analysis. Available research on extremophilic polysaccharides which reported a cryoprotective effect is currently insufficient to derive statistical trends. Therefore, on the basis of a natural selection hypothesis, an assumption of i.e. cold-adapted microorganisms being effective cryoprotectants was adopted to allow for extrapolatory insights and analysis by contrast. The final database (`XPOL-DB`) is composed of various extremophiles, contrasted with mesophilic data, and fed with a database of monomer properties (`MonomerDB`) to enable computation of non-organic parameters, not scraped from the literature (MLAP: monomeric linearly-additive properties). (b) Representation of each extremophilic type for the generation of critical insights. (c) Chronological span of publications for each extremophilic type, reflecting research interest over the years. (d) Parameter completeness plot for each extremophilic type.

**Figure 4:**
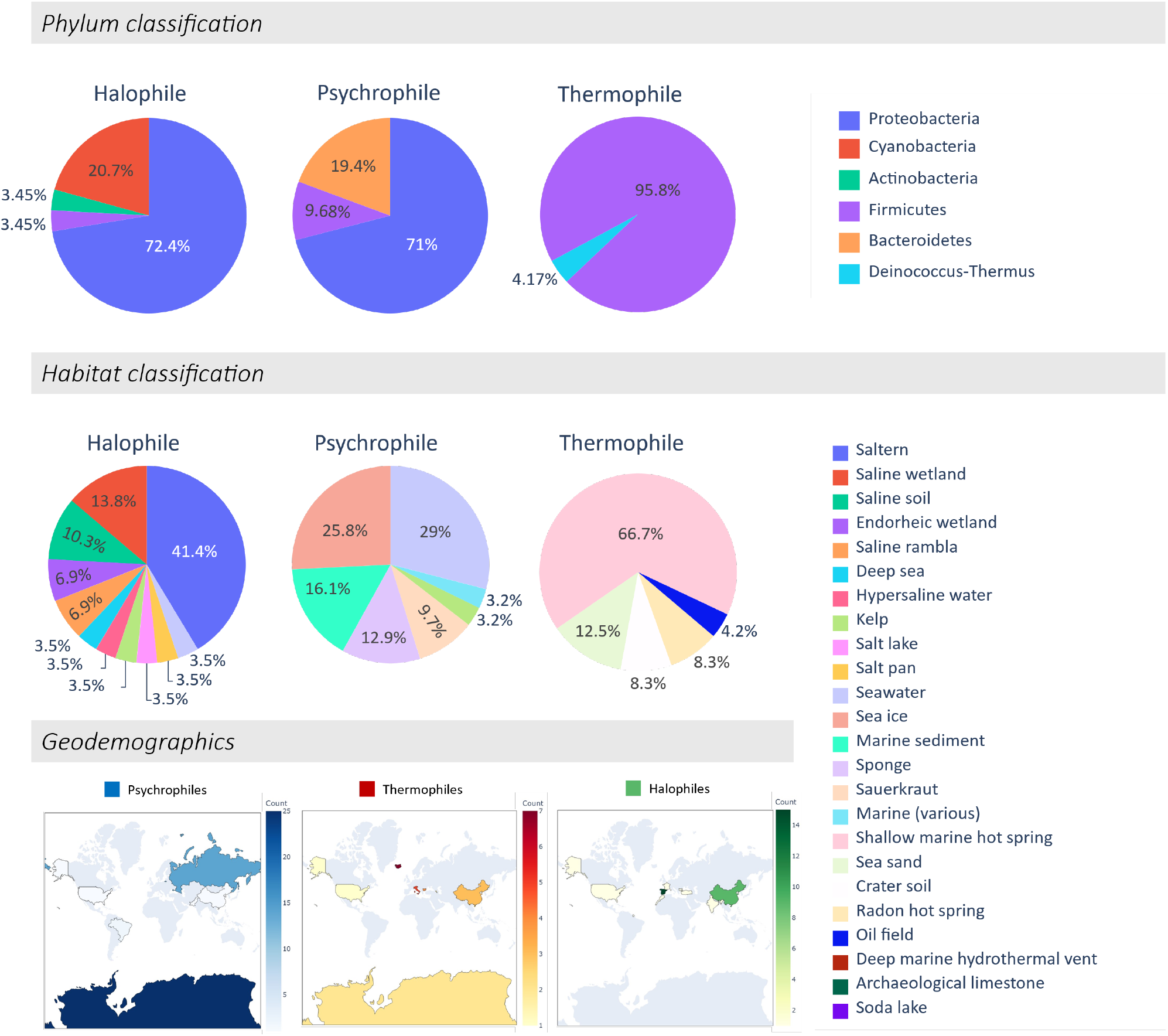
Phylum, habitational and geodemographic distribution of all reported extremophilic hosts reported in `XPOL-DB`, grouped by extremophilic type. For visual clarity, only the three most prevalent extremophilic classes are shown.

#### Habitational markers

The robust nature of extremophiles resulted in a highly diversified habitat footprint. A total of 34 different habitats were identified, some of which biologically shared between different extremophilic types. Psychrophiles were predominantly found in a wide range of cold geofeatures, such as seawater (29% incidence), sea ice (25.8%), marine sediments (16.1%), underwater sponges (12.9%), kelp (3.2%) and deep sea (3.2%), mostly located near the Arctic or Antarctic regions. Halophiles mostly inhabit highly saline geofeatures in equatorial regions, such as salterns (41.4%), saline wetlands (13.8%), saline soil (10.3%), endorheic wetland (6.9%) and saline rambla (6.9%), but they can also be found in psychrophilic habitats, such as marine and deep sea geofeatures with slightly higher salinities, such as deep sea (3.5%), hypersaline water (3.5%) and salt lakes (3.5%). By contrast, thermophiles are exclusive to geofeatures which can either be found in equatorial or polar regions but possess high local temperatures. They can be marine geofeatures regardless of depth, such as shallow marine hot springs (66.7%), radon hot springs (8.3%) or deep-sea hydrothermal vents (3.5%); or dry, such as sea sand (12.5%), crater soil (8.3%) or oil fields (4.2%). Alkaliphiles were mostly found in archaeological sites that contain dry limestone, while hybrid extremophiles prefer wet habitats. Halothermophiles could also be found in shallow marine hot springs where the nutrient composition is rich in salts, while haloalkaliphiles exclusively inhabit salt and soda lakes.

#### Optimal growth & EPS production

The optimal bioreactor cultivation parameters of each extremophilic type for achieving maximal EPS production are summarized in Table 3. There are evident relationships between extremophilic type and main external stressor. Psychrophiles show the lowest optimal average temperature for growth and EPS production (14.9 °C) while thermophiles (45.9 °C) and halothermophiles (54.2 °C) dominate the higher end of the spectrum. Halophiles show the highest salt tolerance (6.9%) and alkaliphiles can thrive in pH 10.4 compared to an average pH 7–7.6 for most species. Psychrophiles require significantly higher carbon source content (C%) in the cultivation medium for EPS secretion across extremophilic types, while nitrogen availability (N%) has a negligible variation. The increased cultivation time necessary for psychrophiles to obtain EPS yields similar to thermophiles has been associated with reduced metabolism due to operation at lower temperatures, but psychrophiles leverage lower temperatures as metabolic optima, thus revealing EPS productivity to be almost 4-fold higher than expected, with a drastic increase in specific yield (2.07 g EPS/g cell dry weight). Figure 5 represents the expected biological ecosystem survivability range for each extremophilic class, a perspective that better demonstrates the biological limits of life than optimal bioreactor conditions in a controlled setting. First, habitat temperature has a narrower tolerance range than pH or salinity. With the exception of psychrophiles, where rare evidence of growth at –40, –50 and –60 °C has been reported, the temperature intervals show rigid boundaries for most extremophilic classes. Con-versely, average pH is observed to be 7.0–7.6 for all classes except alkaliphiles, as they tolerate and thrive in non-physiological pH values. Several psychrophiles have shown optimal growth and production at pH 4–6 and 8–11, halophiles show the same degree of tolerance but at relatively lower acidity, and only mesophiles demonstrate a rigid intolerance towards non-physiological deviations, with a narrow pH regime between 6–8, as expected from non-extremophilic strains. Moreover, all extremophiles show flexible tolerance towards habitat salinity with the exception of mesophiles. Halophilic strains excel at high salinity (up to 30%), but do not necessarily require it for survival, with several accounts of optimal growth between 0–2.5%. Halophiles, halothermophiles and haloalkaliphiles are the only types capable of consistently sustaining salinities above 5%. Psychrophiles are an exception: although they prefer salt contents between 0–5%, some strains have shown tolerance up to 17%. These growth and production metrics provide valuable predictive ballpark values whenever the maximal EPS production conditions are unknown for a specific extremophilic microorganism, drastically simplifying design of experiment strategies.

**Table 3:** Average optimal growth metrics associated with maximal EPS productivity for each extremophilic microorganism type. Parametric ranges (min–max) are indicated by a Δ symbol. Quantitative carbon (C%) and nitrogen (N%) amounts refer to the w/v concentration used in the cultivation medium for each element (most often e.g., glucose for carbon feed and peptone for nitrogen availability). EPS_prod_ refers to maximal EPS productivity. HA: haloalkaliphile; HT: halothermophile. All data sources used for this data aggregation are documented in Table *SI*.*1*, or available online.

| Parameter | Psychrophile | Halophile | Thermophile | Mesophile | Alkaliphile | HA | HT |
| --- | --- | --- | --- | --- | --- | --- | --- |
| T<sub>opt</sub> (°C) | 14.9 | 30.1 | 45.9 | 26.2 | 24.8 | 32.3 | 54.2 |
| $\Delta T$ (°C) | –3 – 26 | 15 – 43 | 46 – 72 | 22 – 38 | 14 – 46 | 11 – 39 | 27 – 65 |
| pH<sub>opt</sub> | 6.9 | 7.2 | 7.1 | 7.0 | 10.4 | 9.2 | 7.6 |
| $\Delta pH$ | 5.5 – 8.1 | 5.7 – 10 | 6.1 – 9.0 | 6.3 – 8.0 | 10.0 – 12.0 | 7.6 – 10.4 | 5.3 – 8.7 |
| Salinity<sub>opt</sub> (%) | 2.5 | 6.9 | 1.4 | 2.6 | 0.8 | 4.8 | 4.5 |
| ΔSalinity (%) | 1.1 – 8.0 | 1.7 – 18.1 | 0.3 – 5.0 | 1.4 – 4.6 | 0.0 – 9.0 | 1.6 – 19.8 | 1.9 – 9.5 |
| C% | 2.2 | 1.4 | 1.0 | 2.7 | 1.5 | 1.3 | 1.6 |
| N% | 0.7 | 0.6 | 0.3 | 0.5 | 0.9 | 0.3 | 0.4 |
| C/N ratio | 5.6 | 8.7 | 3.0 | 6.2 | 2.3 | 6.0 | 2.0 |
| Stirring (rpm) | 141.6 | 97.9 | 244.8 | 509.8 | 120.0 | 150.0 | 155.0 |
| Aeration (v.v.m) | 0.2 | 0.6 | 0.2 | 0.3 | n/a | n/a | 1.0 |
| Time (h) | 117.3 | 181.7 | 28.4 | 56.5 | 114.0 | 88.0 | 60.0 |
| EPS<sub>yield</sub> (g/L) | 1.22 | 3.69 | 1.36 | 6.43 | n/a | n/a | 0.49 |
| EPS<sub>prod</sub> (g/L·d) | 0.45 | 0.72 | 0.11 | 2.71 | n/a | n/a | 0.19 |
| Specific yield (g<sub>EPS</sub>/g<sub>CDW</sub>) | 2.07 | 0.85 | 0.14 | 0.71 | n/a | 2.06 | n/a |

**Figure 5:**
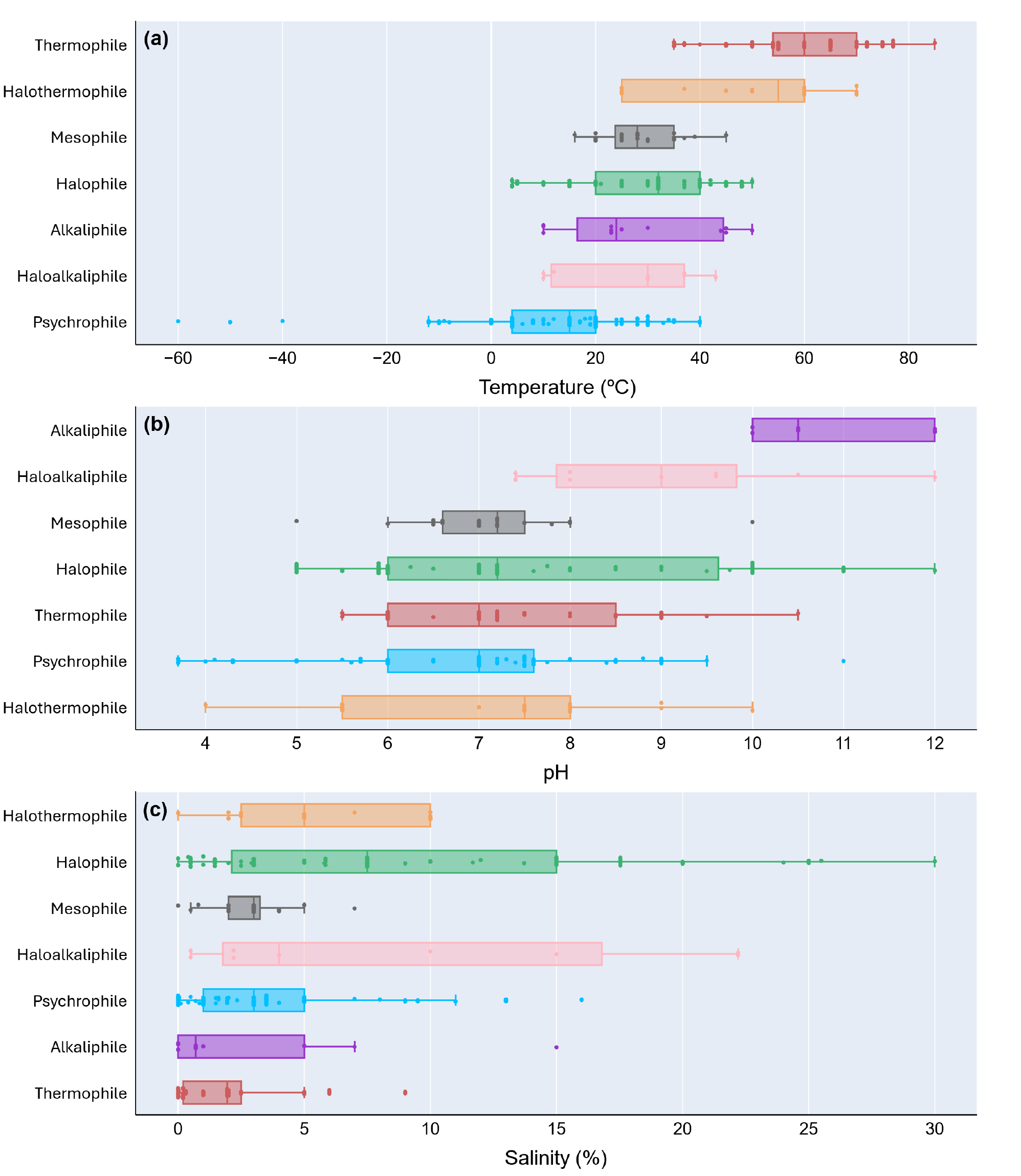
Temperature, pH and salinity spectra for all extremophilic classes. Legend: 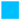 Psychrophile, 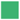 Halophile, 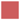Thermophile, 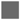 Mesophile, 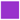 Alkaliphile, 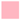 Haloalkaliphile, 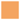 Halothermophile.

#### The psychro-halophilic adaptive similarity

A particular psychrophile, *Colwellia psycherythraea* 34H, arose significant interest. The 34H strain produces an alanine-rich EPS and a threonine-rich CPS (see Figure 12 in further compositional analysis), both branched structures possessing a characteristic pseudohelicoidal conformation similar to the type I AFP (Figure 2), with proven IRI activity and cryoprotective effect [26, 91] due to their amino acid subunits satisfying the hydrophilic-hydrophobic balance previously discussed. Strain 34H is also obligate psychrophile with preferential growth at 16% salinity, which suggests a dual defense mechanism against both stressors. Contrary to psychrophiles and thermophiles being polar opposites of temperature-based stressors, this observation suggests that psychrophiles and halophiles, although arising from different stressors, may share a common survival mechanism. From a physical standpoint, ice growth and ionic gradients are both diffusive mechanisms that rely on cell membrane osmoregulation. In extreme conditions, both phenomena create a physical membrane destabilization effect that generates hydrostatic pressure. At the later stages of ice growth, the incremental frozen fraction leading to solute supersaturation establishes a microenvironment dynamically equivalent to brine channels [2, 16], characterized by an increased concentration of solutes (i.e. salts). Thus, the halophilic mechanism for salt tolerance might strongly correlate with the psychrophilic mechanism for freezing tolerance. Revisiting habitat temperature for each extremophilic class with greater granularity (Figure 6), psychrophiles and halophiles show very similar temperature regimes and tolerance. Moreover, both extremophilic types can co-inhabit sea water, deep sea and submarine kelp. The fact that thermophiles and halothermophiles strictly inhabit high temperature regions, alkaliphiles find their unique habitats in regions where stone can accumulate specific oxidizing metals, such as archaeological limestone, and haloalkaliphiles exclusively populate salt and soda lakes (pH 12), further supports the observation that psychrophiles and halophiles show the greatest adaptive pressure similarity relative to any other pair combination of extremophilic types, and the highest degree of structure-function relationship convergence relative to all other pairings.

**Figure 6:**
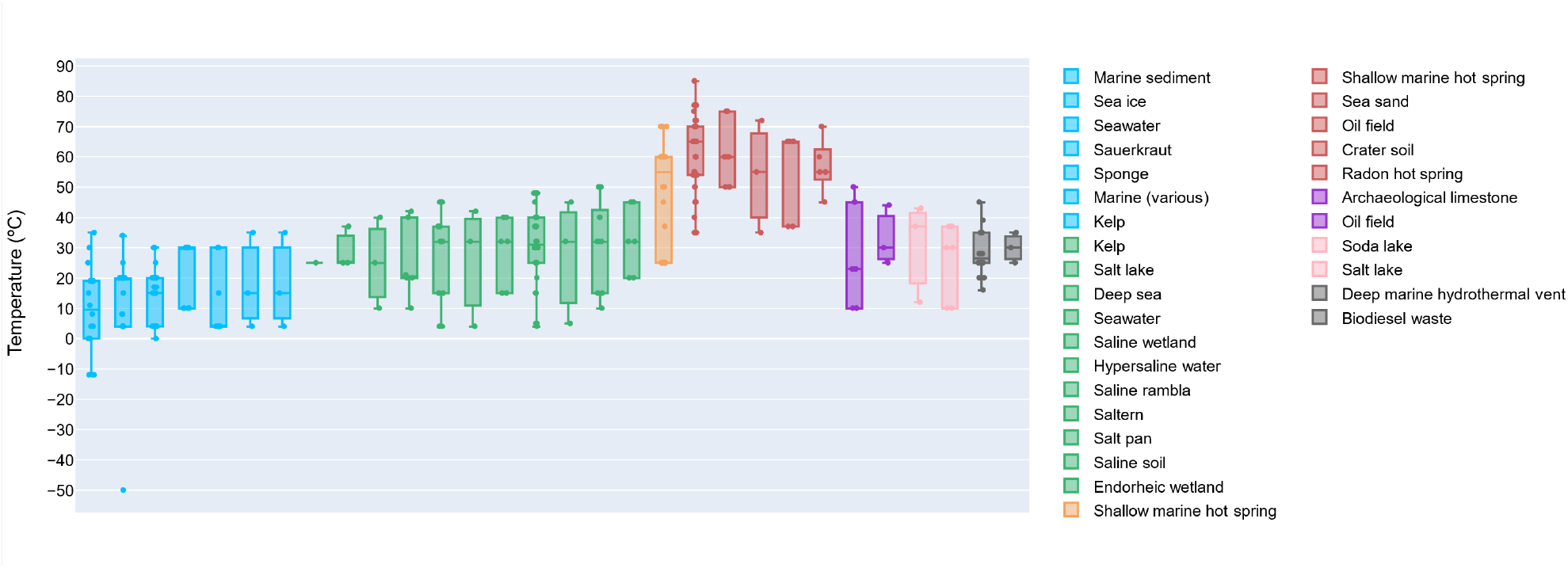
Temperature ranges for each habitat geofeature. Legend: 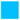 Psychrophile, 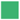 Halophile, 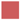 Thermophile, 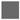 Mesophile, 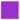 Alkaliphile, 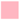 Haloalkaliphile, 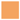 Halothermophile.

#### Molecular weight

Most produced EPS structures range between 10^3^–10^4^ kDa, except halophiles and psychrophiles which average at higher M_w_ distributions (Figure 7). Psychrophilic EPS show M_w_ ranging from simple 6.5 kDa chains up to immensely com-plex structures 300 MDa in size, with an average 16.2±56.8 MDa (N=38). Halophiles show an average M_w_ 1.7±3.6 MDa (N=29), followed by halothermophiles (1.4±2.2 MDa, N=5), mesophiles (1.1±1.6 MDa, N=15) and alkaliphiles (1.1±1.8 MDa, N=4). Thermophilic polysaccharides hold both the lowest average M_w_ (0.59±0.72 MDa, N=19) and the narrowest size distribution of all types. M_w_ is often correlated with structural complexity. Greater prevalence of chain branching at high M_w_ regimes leads to increased solution viscosity due to an acquired resistance to flow [108]. In the context of cryobiology, an increased viscosity often correlates with ice growth inhibition due to hampered solution dynamics required for molecular aggregation and hydrogen bond directionality during nucleation and growth. Hence, the observation that psychrophiles possess the highest M_w_ observed for all extremophilic types is expected, and most likely contributes to freezing avoidance by reduced kinetic rates and increased mechanical resistance in the EPS architecture. Once again, both psychrophiles and halophiles share a structure-function relationship similarity in producing polysaccharides with above-average molecular weight, with certain halophilic polysaccharides reaching M_w_ values ca. 2–19 MDa. This insight is further supportive of an increased M_w_ being contributive towards inhibiting solution kinetics, to osmoregulate the passive diffusion of supersaturated salt media through the host microenvironment, in order to alleviate osmotic shock.

**Figure 7:**
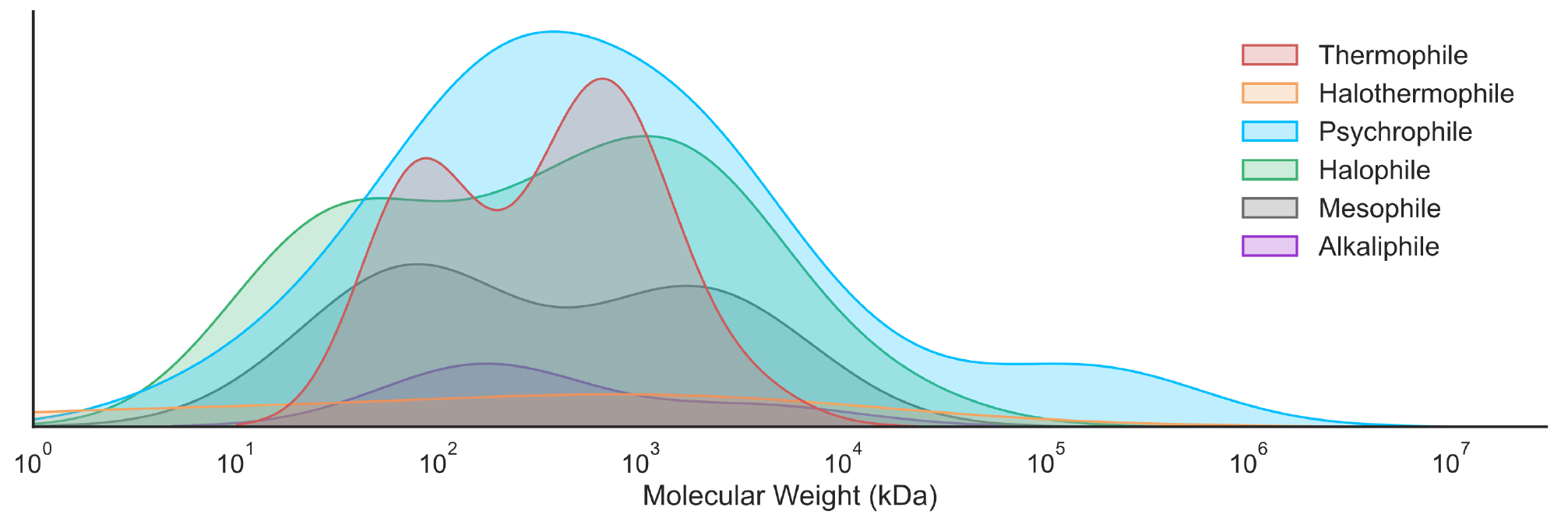
Molecular weight distributions of extremophilic EPS, grouped by class. Psychrophilic: N=38; Halophilic: N=29; Halothermophilic: N=5; Mesophilic: N=15; Alkaliphilic: N=4; Thermophilic: N=19. The y-axis is an arbitrary scale of M_w_ incidence.

#### Compositional fingerprinting

The core structural feature of an extremophilic polysaccharide is its monomer composition. The structural repeating unit (SRU) contains all chemical information that will determine net formal charge, structural conformation, physical properties and biological function, by liaison of their monosaccharide content, respective linkage types and degree of branching. Hence, a complete chemical characterization of their monomer content was considered a foundational requirement in `XPOL-DB` inputs. A frequency analysis of the monosaccharide composition of all 145 EPS, out of which 60 are psychrophilic, was carried out and is summarized in Figure 8. The neutral monomers glucose (Glc), galactose (Gal) and mannose (Man) possess the highest incidence among all monosaccharides. Glc is present in 77.4% of extremophilic polysaccharides (67.8% in those psychrophilic), followed by Gal (62.9% globally, 67.8% psychrophilic) and Man (67.1% and 52.5%, respectively). Comparing global and psychrophilic monomer incidences enables discerning which monosaccharides are significantly most prominent in cold-adapted polysaccharide structures. Overall, psychrophilic polysaccharides are richer in polar monomers, containing GalNAc (+12%), GalA (10.9%), GlcNAc (+9.3%), GlcA (+4.1%), QuiNAc (+4%), GlcN (+3.6%) and Kdo (+2.7%) in their structural compositions, than the global average. The common denominator for all these monomers is a net formal charge, expressed in one of three forms:

**Figure 8:**
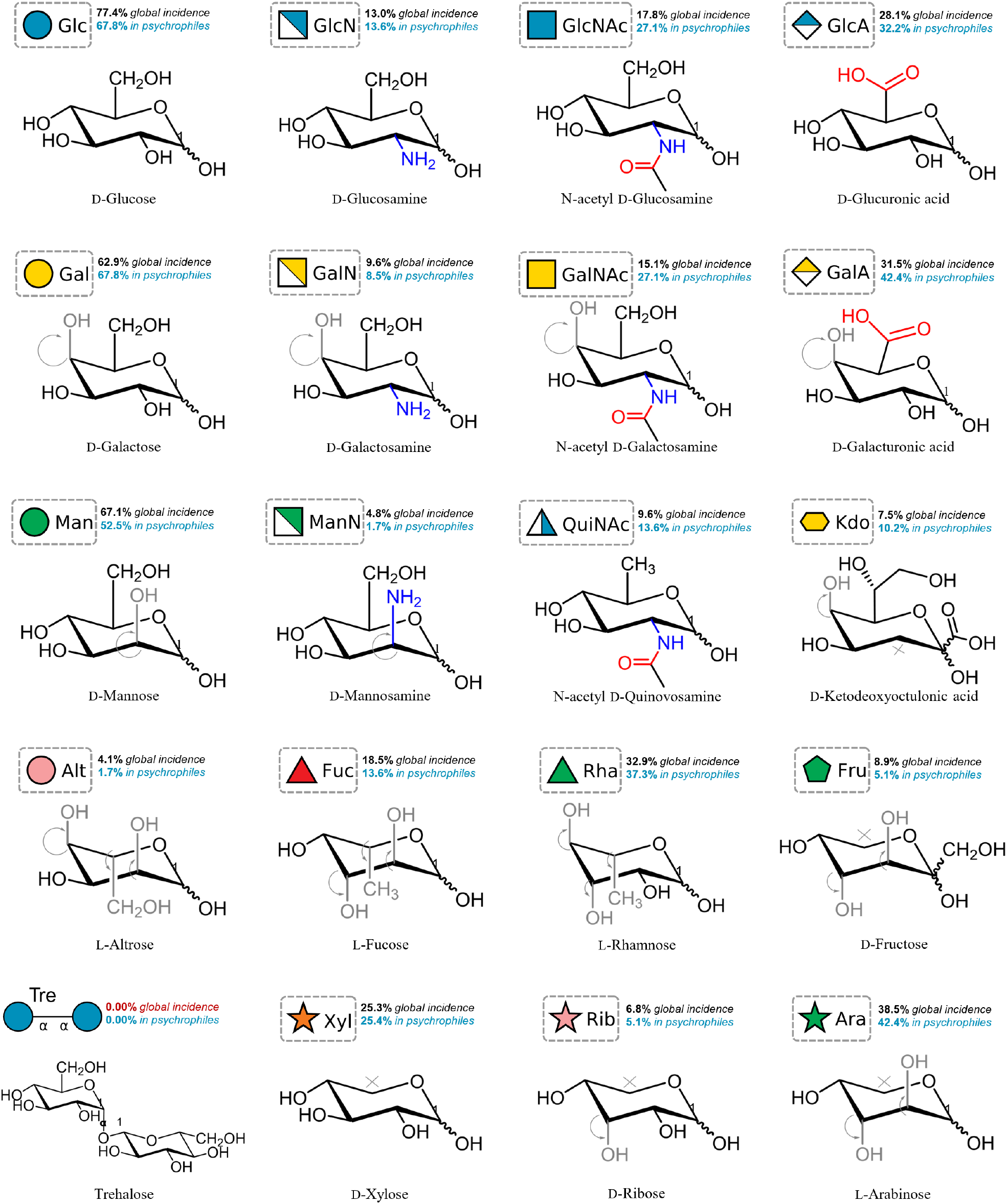
Monomer frequency analysis across extremophilic types. The global and psychrophile-only prevalences for each of the top 20 most frequent monomers in the database are reported, accompanied by the standardized Symbol Nomenclature For Glycans (SNFG) [109]. An enhanced psychrophile-only prevalence relative to the global incidence is suggestive of a significant contribution of a given monomer towards cold adaptation. From left to right, each column represents neutral, cationic, zwitterionic and anionic monomers, respectively. The last two rows, with the exception of altrose and trehalose (absent in polysaccharides), cluster monomers which do not contain a hydroxymethyl moiety on C_6_ or no C_6_ at all.

1. anionic carboxyl groups at C_6_, axially distributed (e.g. uronic acids),
2. cationic amine groups at C_2_, equatorially distributed (e.g. hexosamines),
3. zwitterionic amide groups at C_2_, equatorially distributed (e.g. N-acetylated sugars).

All other neutral monosaccharide forms appear mostly underrepresented in psychrophiles, possibly due to a lack of formal charge not being beneficial towards function. Monomers of L-configuration such as the methylated Rha (+4.4%) and the non-methylated Ara (3.9%) are also present. Fucose (Fuc) is an intriguing L-monomer, because it appears to be exclusive to psychrophiles and halophiles, but not thermophiles. Trehalose (Tre), a disaccharide highly acclaimed for its cryoprotective potential [110], is never present in polysaccharide structures, prompting further investigation into whether its absence is due to structural inadequacy, lack of genetic glycosyl transferases specific for trehalose, or a non-emergence of cryoprotective potential in a polymeric structure. The creation of compositional fingerprints for each extremophile, and comparison of the psychrophilic fingerprint with that of other classes allows to discern patterns of monomer occurrence. By the recurring postulation of Darwinian selective adaptation having a phenotypical expression in the structural traits of molecular defense mechanisms, then the overexpression of certain monomers in particularly exclusive classes yields valuable insight into how their properties may contribute to expressing particular traits at a macroscale. Figure 9 shows comparative contrasts between the relevant compositional fingerprints obtained for each extremophilic class. In general, the neutral monomers Glc, Gal and Man are of ubiquitous occurrence amongst all types. This suggests a non-specific structural linker role rather than specific bioactive adaptation towards a particular functionality (Figure 9a). Man is present in higher quantities in extremophilic strains. In mesophiles, Gal and Glc dominate instead. Although they occur as core monomers, that does not exclude a possible influence in enabling particular structural conformations in-between bioactive moieties, such as is the case of the neutral Ala amino acids in the type I AFP previously described, whereas they did not participate in ice-binding but conferred helical flexibility to polar binding Thr for better compliance to the ice surface curvature. Figure 9b shows the standalone compositional fingerprint of psychrophilic EPS, which contains the greatest compositional diversity, with a total of 22 different constitutional monomers. Their distinguishing feature is the predominance of polar monomers, highlighted by a monomer content equal or higher than the shown extremophilic average (black dot), namely GlcN, GlcA, GalA, Kdo, GlcNAc, GalNAc, Rha, Fuc and QuiNAc, with some neutral monomer exceptions (Rha, Fuc), as previously pointed out. The compositional superposition of psychrophilic and halophilic fingerprints in Figure 9c shows that, with the exception of very low incident GalN and Rib, both classes share not only the same monomer diversity but a very similar fingerprint relative to other pairings. The less visible difference lies in an accentuated amount of neutral monomers in halophiles, which is compensated in psychrophiles by an accentuated amount of anionic and zwitterionic monomers. However, the strong profile superposition reveals that there is a high similarity between psychrophilic and halophilic compositions, reinforcing the previous inference of a shared mechanistic denominator between their specific external stressors. Conversely, Figure 9d highlights mechanistically opposed adaptation trends in the temperature domain, i.e. between psychrophilic and thermophilic structures. Thermophilic polysaccharides not only are less diverse in monomer composition, but strongly express arabinose (Ara) and xylose (Xyl) in their structures, alongside the rarer altrose (Alt) and ribose (Rib), rather than anionic or zwitterionic species, highlighting the influence of the latter in cold adaptation but not heat. Figure 9e further emphasizes this fingerprint dualism between psychrophiles and halophiles; and thermophiles. The particular observation that certain neutral monomers (Rha and Fuc) but not all have a strong incidence in halophilic and psychrophilic polysaccharides but not thermophiles, despite their neutral character, prompted further investigation. In a subsequent study, a large bio-based polysaccharide screening revealed that fucose-rich, polyanionic polysaccharides drastically enhance the survival rate of cryopreserved cells, when compared to non-fucose counterparts; and their contributive mechanism is not related to electrostatic disruption but rather conformationally enabling (analog to type I AFP Ala residues) and potential membrane stabilization effects [62].

**Figure 9:**
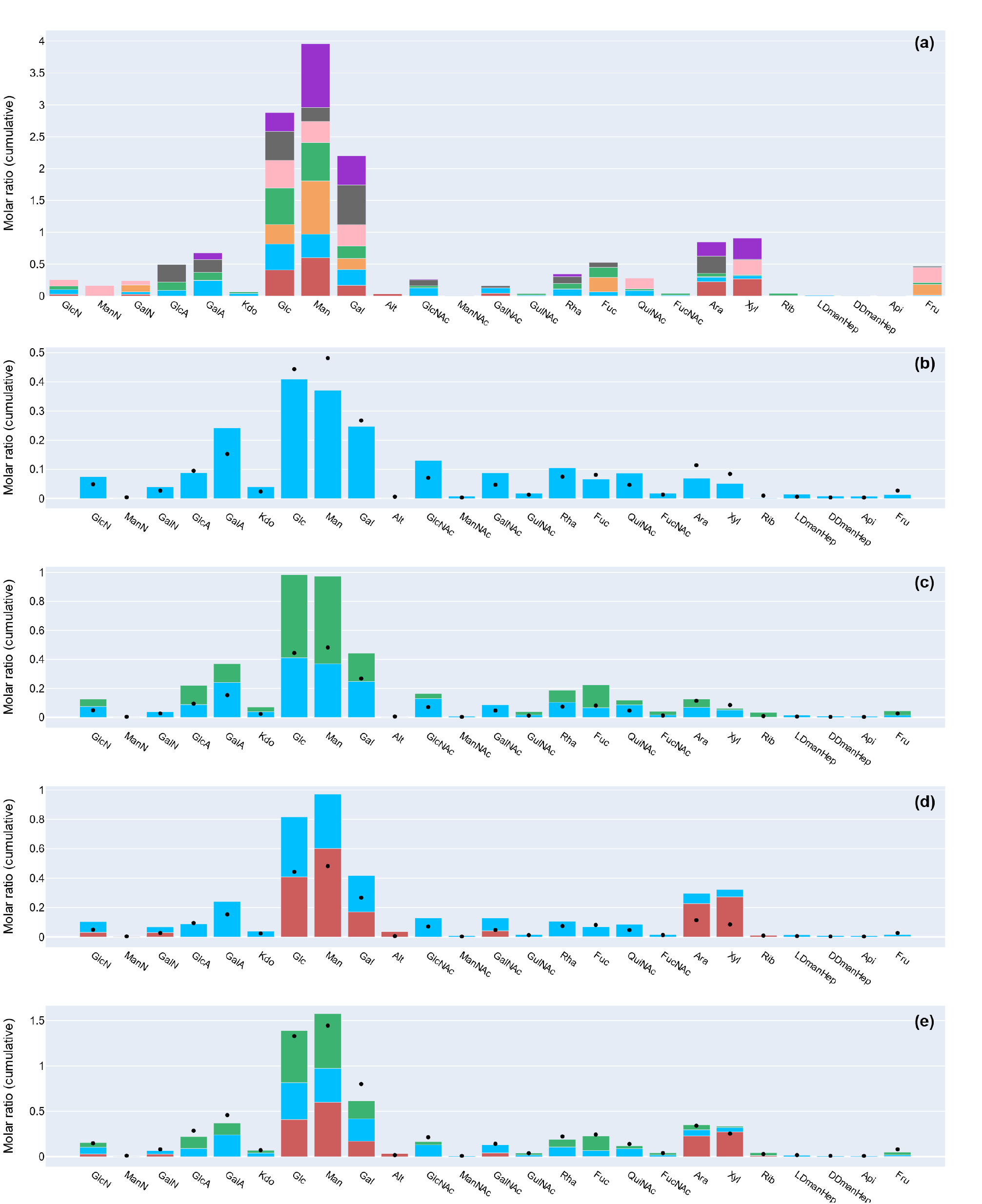
Average compositional fingerprint of each extremophilic type, revealing distinct monomer prevalences indicative of specific functionality. Panel (a) presents a general overview, while panels (b–e) show relevant comparisons between psychrophiles, halophiles and thermophiles. The black dot represents the weighted average molar ratio for a given monomer for the extremophilic types plotted. Legend: 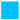 Psychrophile, 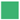 Halophile, 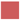 Thermophile, 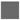 Mesophile, 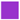 Alkaliphile, 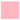 Haloalkaliphile, 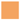 Halothermophile.

#### Polarity maps

Compositional fingerprinting is a very powerful visualization tool because it hinted that (i) certain monomers are intentionally embedded in polysaccharide structures because they provide a chemical feature beneficial towards dealing with the external stressors that exclusively arise in cold and highly saline habitats; and (ii) some halophilic polysaccharides may indeed be cryoprotectant. The strong significance of net formal charge in being a differentiating feature between psychrophilic and thermophilic EPS prompted the representation of compositional fingerprints as a function of polarity (Figure 10). By close inspection of the red and blue shadowed regions in the radial charts, which represent charged monomer species, there is renewed corroboration that halophilic EPS may express biological traits similar to psychrophilic EPS due to similar polarity maps. In contrast, thermophilic EPS are mostly neutral structures (beige). Thus, a chemical preference exists for polyanionicity towards water bond interaction and successful ice growth disruption in cold-adapted environments.

**Figure 10:**
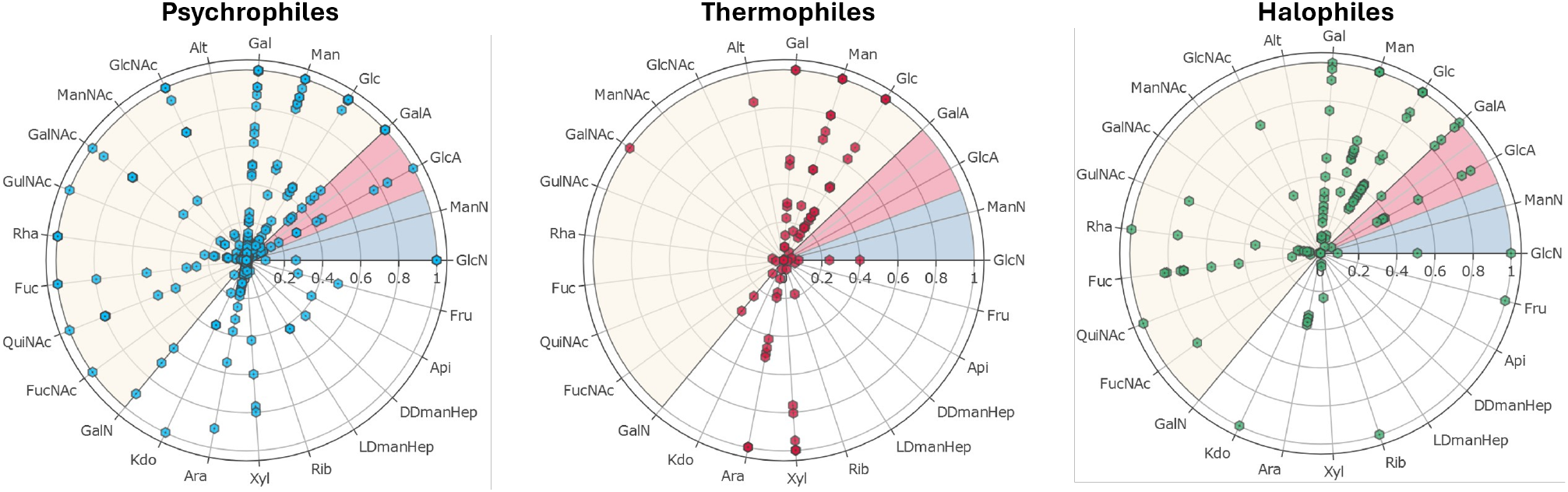
Radial representation of the compositional class-level fingerprints for psychrophilic, thermophilic, and halophilic polysaccharides. The beige, red, and blue shaded regions represent neutral, anionic, and cationic monomers, respectively. The *x*-axis represents the normalized molar content of each monomer in EPS structure.

### Biological function

Extremophilic EPS were also probed for their different biological functionalities. Figure 11 shows a correlational matrix that maps how each particular biological function reported in the literature can be associated with an extremophilic type, or its characteristic external stressor. Three major correlation clusters can be observed near the identity line (black). First, the leftmost-top cluster aggregates functional forms of avoiding ROS-based damage, such as antioxidant, antitumoral and antiradiation properties (*R*^2^ = 0.1–0.4). Second, the centermost cluster appears related to kinetic and structural properties (*R*^2^ = 0.2–0.5). Heavy metal binding (HMB), emulsifying, swelling and gelling properties are associated to interchain non-covalent binding, leading to structural transformation of the polysaccharide architecture, which may alter viscosity and diffusional kinetics of the surrounding and confined solvent, thus mediating passive diffusion. In a subsequent study, these were later corroborated to be critical aspects of cryoprotective polysaccharides [63]. Third, the rightmost-bottom cluster relates to internal host regulation responses, and includes immunoregulatory, antibacterial, antiviral, antiapoptotic, anticoagulant and anti-inflammatory functions (*R*^2^ = 0.1–0.5). The first and third clusters also appear co-correlated, which indicates that radiochemical and biological triggers to the immune system are naturally exogenous, and thus present an acute stressor, to which biofilm polysaccharides have acquired protective strategies against. For the subset of cryoprotective polysaccharides, only 33 accounts of quantitative cryoprotective activity were reported, and the reporting methodology varies extensively. Most data was obtained from red blood cell (RBC) cryopreservation (36%), host self-preservation (12%),

**Figure 11:**
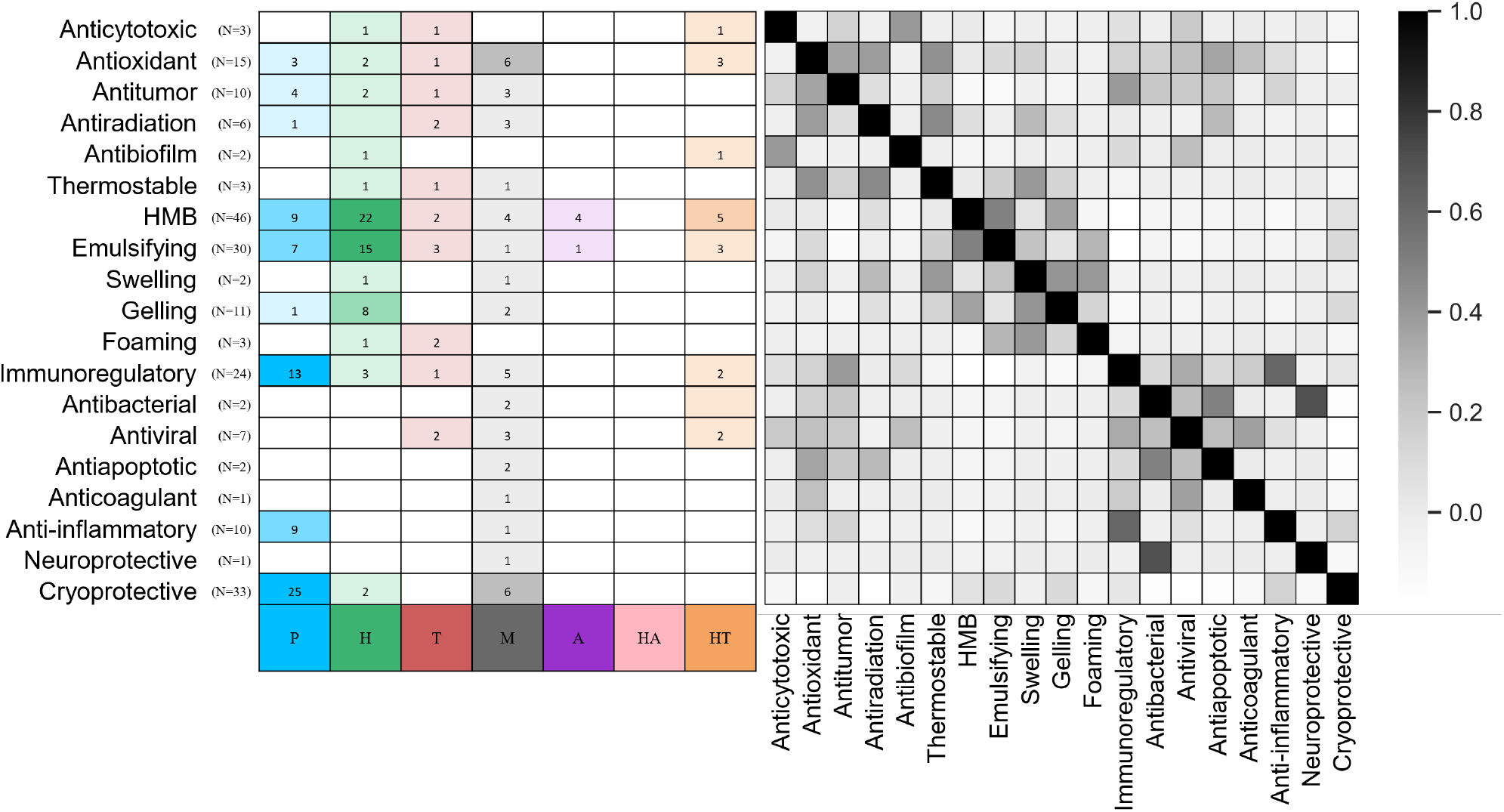
Biological functionality matrix, showing observation frequency by extremophilic class (left) and the respective correlation heatmap (right). The color gradient refers to *R*^2^ correlation data. P: psychrophile; H: halophile; T: thermophile; M: mesophile; A: alkaliphile; HA: haloalkaliphile; HT: halothermophile.

*E. coli* vector freezing (9%) or reproductive cell post-thaw fitness after cryopreservation (9%), such as sperm (6%) or oocytes (3%). Most procedures involve slow freezing using cooling rates in the 1–3 °C/min range (58%) or fast freezing with liquid N_2_ (42%), by employing either a single freeze-thaw cycle (67%) or up to four freeze-thaw cycles (33%), held for cryostorage periods between 24 h and one week. The mechanisms of action by which polysaccharides exerted their cryoprotective effect are often underreported. Of those demonstrating some form of cryoprotective outcome, 70% were observed to enhance biological survival, using different biomarkers for validation, such as leukocyte count (27%), post-thaw metabolic activity (9%), viable cell count (3%) or post-thaw growth rate comparison (3%). Moreover, IRI activity and ice growth inhibition were both reported 21% of the time, nucleation effects 9%, anti-nucleation effects 3% and dynamic ice shaping 3%, with the most dominant being thermal hysteresis (TH), for a total of 39%. The fact that the cryoprotective action of polysaccharides has been linked to a strong influence on nucleation phenomenology [60] but remains largely underreported in the literature as a mechanism of action (3–9%) indicates a suboptimal understanding, also largely acknowledged before [111], of polysaccharide cryoprotection mechanisms. Based on Figure 11, extremophilic polysaccharides with potential cryoprotective action show appreciable correlations with HMB, emulsifying, gelling, immunoregulatory and anti-inflammatory properties. The cryoprotective polysaccharide FucoPol has shown a concerted multifunctionality that spans all these traits [63, 96, 104, 112, 113], further corroborating these findings from `XPOL-DB`. Ultimately, the data suggests that common structural traits between extremophilic polysaccharides result in the expression of (i) multifunctionality, and that (ii) diverse biological functions are associated by common chemical mechanisms, suggesting the existence of a unified understanding that can assist on reverse engineering desired polysaccharide compositions based on intended functionality.

#### Structural Repeating Units (SRU)

In proteins, the peptide linkage contains a single N-terminal and C-terminal pair and defined angular constraints given by the Ramachandran plot, but protein structure is largely defined by amino acid composition. In polysaccharides, there is a particular distinctiveness of the glycosidic linkage, the linkage itself being a defining constraint to structure, as several combinations of *α*- or *β*-linkages and anomeric carbon bond (1→ 4, 1→ 6, 1→ 3, etc.) may exist, thus limiting which monomer can link to the next position in the chain. This intricacy in polymerization influences every other aspect of structure, from molecular weight to composition, from low to high-level organization, from the sequence of the structural repeating unit to expressed function. For instance, an *α*(1→ 6)-linked glucan constitutes the highly soluble, flexible, globular starch; but a minute change to *β*(1→4)-linked glucan yields the highly insoluble, rigid, interconnected mesh-like cellulose. Figure 12 represents a subsample of the large cohort of biotechnologically produced extremophilic EPS structures reported in `XPOL-DB`, alongside commonplace commercially used polysaccharides, to provide a contrast between the composition of their SRUs. The difference in SRU complexity between the reported extremophilic EPS in `XPOL-DB` and the commercially sourced mesophilic structures is noticeable. In general, there is a greater monomeric diversity amongst different extremophilic types; and a greater incidence of terminal uronic acids, acetylation or amino acid residues than in mesophiles, which may contain polar species but in internal domains of their structure. However, some similarities exist, such as the psychrophilic SM20310 or SM9913 with non-psychrophilic dextran or starch, all of these being glucans. There is a particular incidence of negatively charged monomers and branching in psychrophilic polysaccharides (e.g. *Colwellia psycherythraea* 34H), compared to the linear N-acetylated thermophilic galactan TA-1 from *Thermus aquaticus* YT-1. Water-insoluble polysaccharides express mostly simplistic, linear, neutral structures.

**Figure 12:**
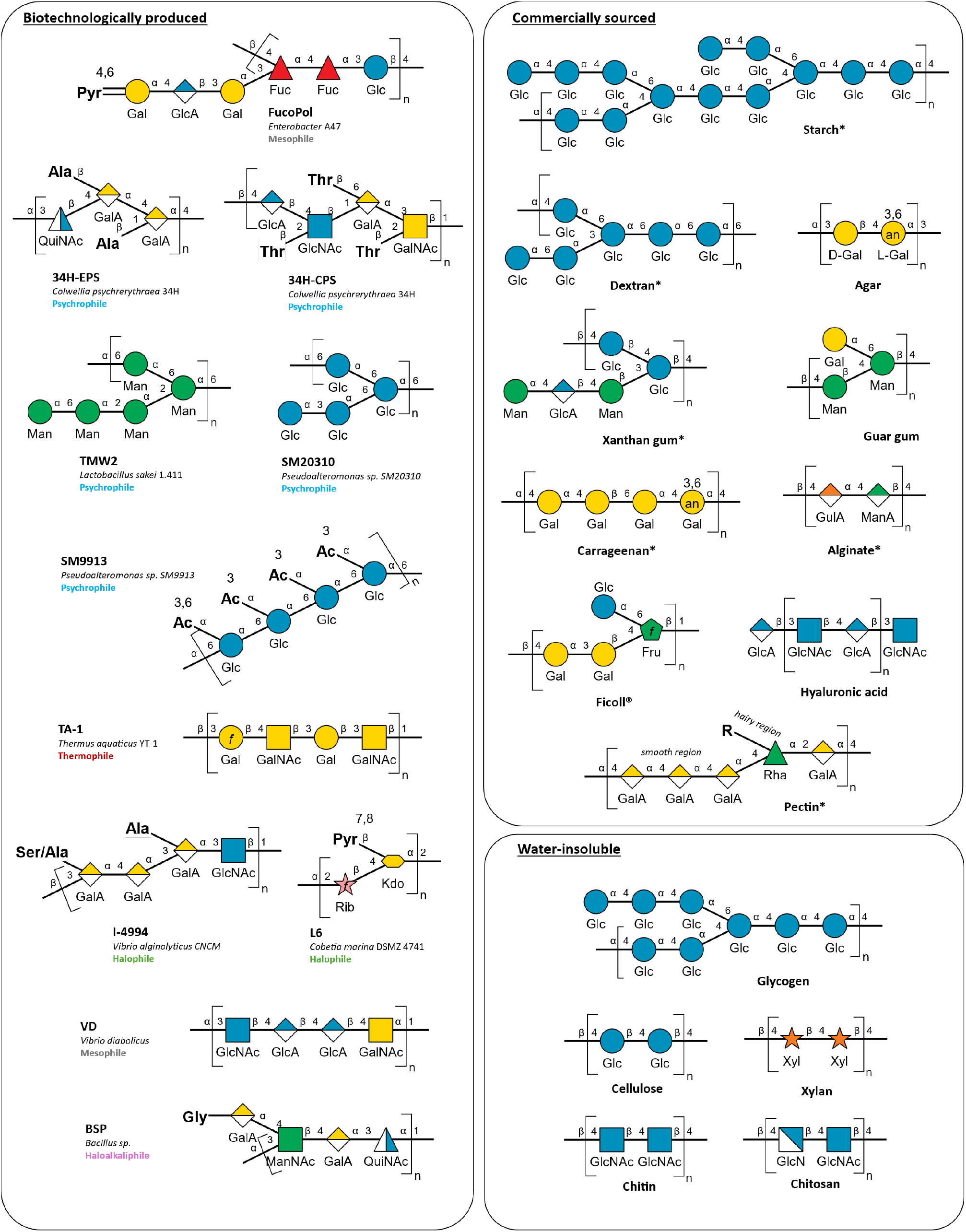
Structural repeating unit (SRU) representation of relevant polysaccharides. Some exemplary extremophiles from the `XPOL-DB` database are shown (“biotechnologically produced”). The right panel shows common commercially sourced polysaccharides, and water-insoluble structures, for comparison. Polysaccharides labelled with an asterisk (*) represent the general backbone pattern of a family of structures. The SNFG symbology standard was used [109].

#### Conformational profiling

The fact that the psychrophilic SM20310, starch, glycogen and cellulose may have completely different degrees of solubility and bioactivity despite similar structures suggests that composition alone is not sufficient to justify a perceived property. So, we leveraged the structural reporting in `XPOL-DB` to estimate which structural patterns were most often associated to a given extremophilic stress factor, to inquire how features other than chemical composition could impact their role in stress adaptation. Unlike protein structures which follow a standard reporting of their secondary and tertiary structure, structural reporting in polysaccharides is done loosely. Figure 13 presents a putative structural hierarchy based on several reports. For the extremophilic EPS of known composition and structure, the primary conformation is usually ambiguously reported to be either linear/random coil (30.2% incidence) or branched (21.4%). Psychrophilic polysaccharides mostly present branched backbones (56.7%), unlike thermophiles which produce mostly linear polysaccharides (41.7%). Halophilic polysaccharides are also highly branched (31.3%). In terms of secondary structure, four features have been identified: hyperbranched (16.6%), helix (2.1%), pseudo-helix (1.4%) and hairpin (0.7%) conformers. Rare accounts of four tertiary conformations have been reported, usually reflecting chain flexibility and interchain associations. Polysaccharide backbones of linear nature (e.g. thermophiles) usually form rigid rod-like or flexible worm-like structures from intrachain coiling. Conversely, highly branched polysaccharides (e.g. psychrophiles and halophiles) preferentially organize into entangled mesh-like sheets (non-covalent) or reticulated networks (covalently cross-linked). Polysaccharides have been reported as cationic (2.1%), zwitterionic (2.8%), neutral (11.7%) or anionic (62.8%), the latter present across all extremophilic classes. However, 32.4% of anionic EPS are psychrophilic and 19.3% are halophilic, compared to thermophiles (14.5%) and mesophiles (9.7%).

**Figure 13:**
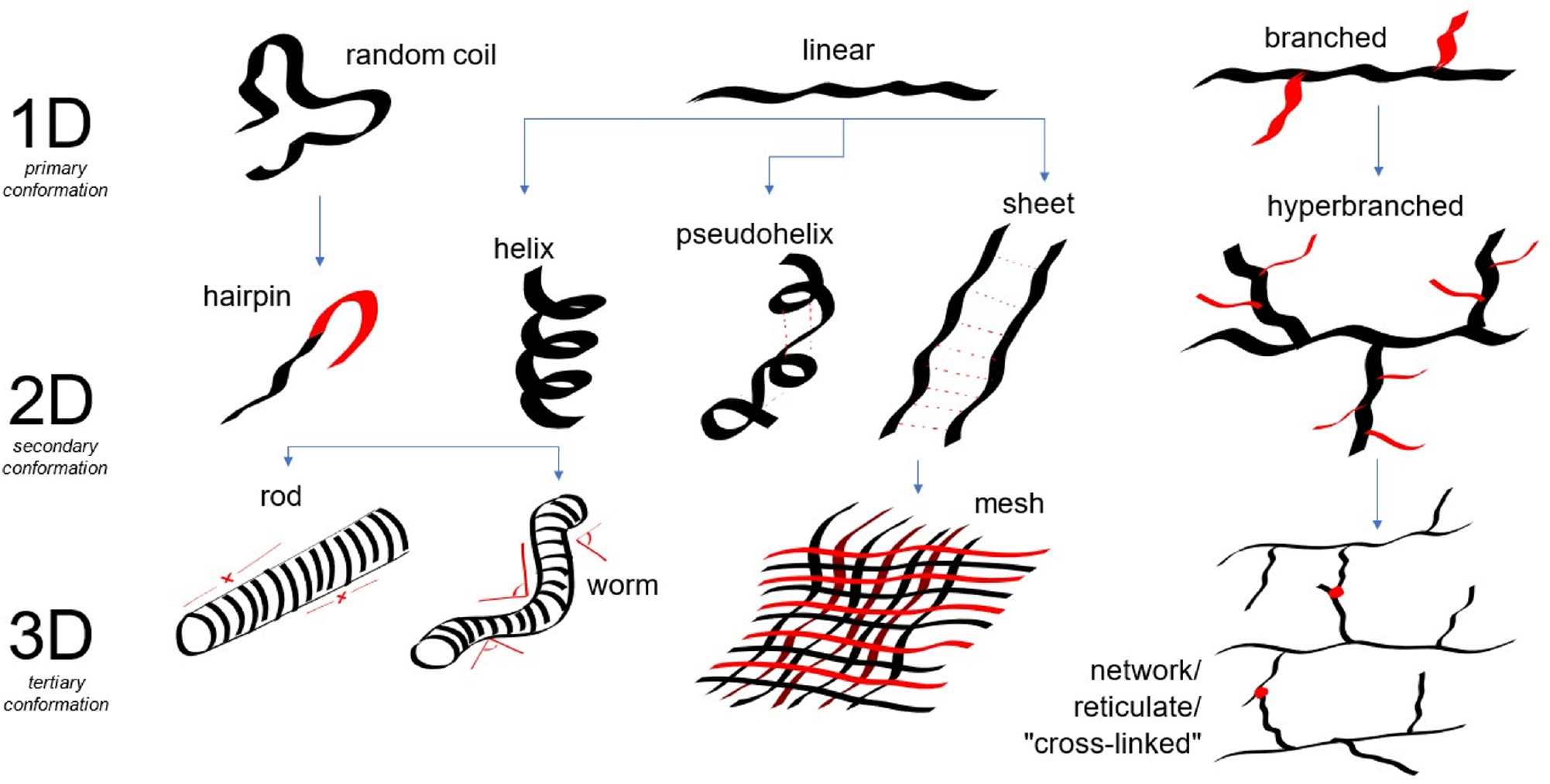
Illustration of the perceived structural hierarchy of the potential primary, secondary and tertiary conformations for the extremophilic EPS reported. Domains drawn in red represent distinctive features for contrasting visual clarity. The blue arrows indicate a parent-child relationship between structures of different degrees of conformation, and were often reported to subsequently lead to one another based on similar spatial re-arrangements and properties.

#### Glycosidic linkage profiling

Given the significance of a different glycosidic linkage having a dramatic effect on the properties of same-composition glu-cans, a complete frequency analysis of the glycosidic linkages present in the known SRUs of extremophilic polysaccharides was mapped (Figure 14). Psychrophilic polysaccharides of known complete structure (N=37) are rich in *α*-linkages (4.2 *α* per SRU) compared to *β*-linkages (2.0 *β* per SRU). Psychrophiles are rich in *α*(1→2, 3, 4) and *β*(1→3, 4, 6) linkages, which indicates that hyperbranching most occurs by *β*-linkages at the C_6_, whilst the backbone is evenly constituted by *α* and *β*. Halophiles present a similar linkage profile, although the bond character at the anomeric carbon is rarely reported. However, the presence of ?(1→6) linkages suggests some degree of branching, as previously confirmed; and is distinctive from thermophiles, very scarce in 1→6-linked bond types, which is validated by their linear structures. The predominance of *α*-linkages is similar for all classes, so an *α*/*β* ratio demonstrated to be more informative than an absolute linkage type count. Psychrophiles and halophiles expressed *α*/*β* ratios of 2.0 *±* 1.2 and 1.6 *±* 1.4, respectively, compared to thermophiles (1.1 *±* 0.4) and mesophiles (1.2 *±* 1.2), with a slightly more balanced bond character. This suggests that a relative predominance of *α*-linkages may be associated with biological functionalities which require chain pliability (e.g. ice binding affinity), while *β*-linkages may contribute predominantly to chain rigidity and stabilization (e.g. thermostable molecules).

**Figure 14:**
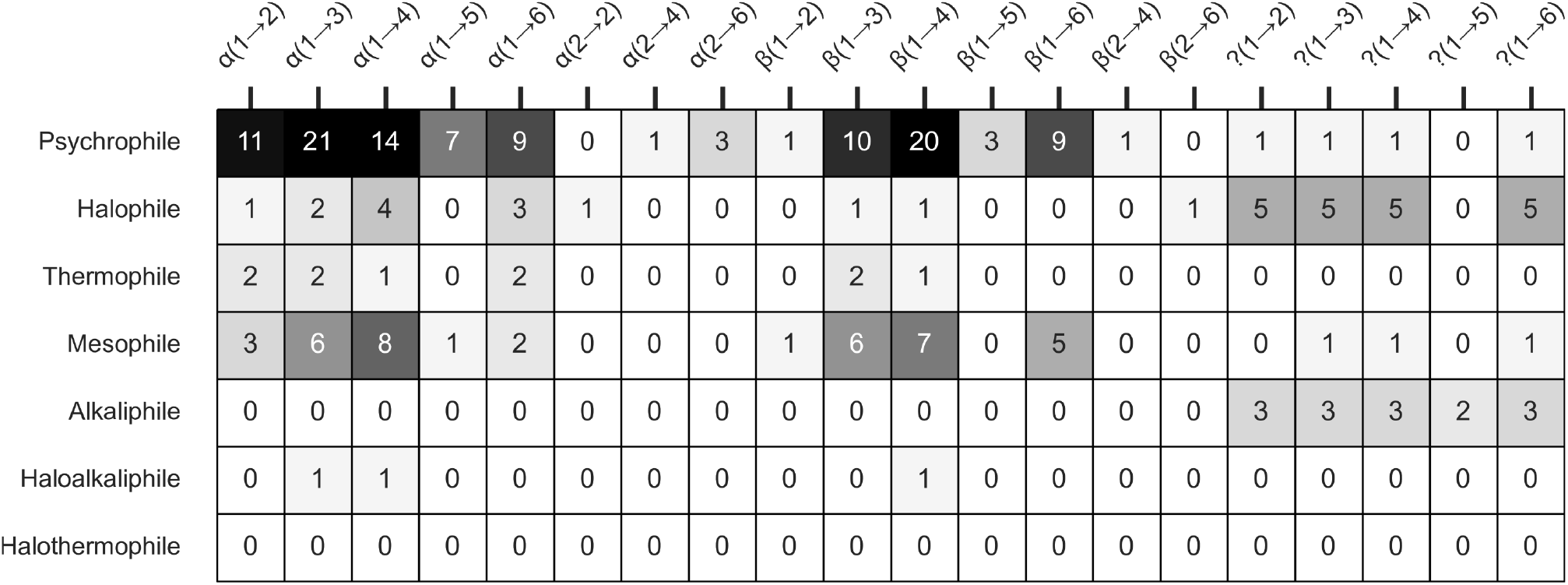
Glycosidic linkage type distribution across extremophilic polysaccharide classes, mapped using a one-hot encoding algorithm. The ?(*x*→*y*) bond symbology refers to an underreported unknown disposition of the anomeric carbon C1 relative to the *n* + 1 sugar monomer in the linkage, but known participating carbon atoms in the linkage.

### Multidimensional Data Modelling

The multidimensional character of `XPOL-DB` allowed to derive new literature-driven insights from sparse structure-function relationship trends, but it does now confer an immediate practicality to its use. In highly complex databases, an extensive reduction of the universe of relevant attributes for manual lab experimentation far outweighs any attempts at automating the analysis of the whole parametric universe. This is because the variables that maximize impact in outcome are often a fraction of the useful variable space, and focus on those minimizes the time from experimental design to insight gathering. Thus, to further highlight which structure-function traits could have the highest likelihood of indicating a cryoprotective potential, we performed dimensionality reduction techniques and subsequent parametric screening. By filtering for the most physically meaningful attribute correlations, predictive analysis can determine the irreducible complexity required for the expression of function from a small subset of characteristics.

#### Batch correlation analysis & variable filtering

The `XPOL-DB` contained a total of 128 reported attributes for the characterization of each extremophilic EPS. First, we performed a pair-plot batch correlation analysis, in which a correlation factor *R*^2^ was calculated between every combination of existing variable pairs. After discounting self-correlations (*R*^2^ = 1) and matrix reflection duplication across the identity line, a total of 6352 correlations were identified in the database. Naturally, experimentally assessing all these is an unfeasible and largely redundant workload. First, Figure 15a revealed that the number of correlations decreases with increasing *R*^2^ in exponential fashion, and that most correlation pairs are located in the *R*^2^ = 0–0.2 region. Second, Table 4 informed that almost half of all correlations are above *R*^2^ = 0.1 (48.3%, 3070 correlations). Third, Figure 15b indicated that some observational bias is present in the database: with a decrease in the number of correlations (i.e. higher *R*^2^), the number of collected parameters in the database linearly decreased from *N* = 140 to about *N* = 5. The prediction bands are highly divergent from the average trendline, which indicates data reporting in the literature is not consistent for most variables. Thus, Figure 15c computed a weighed number of correlations, in which the product between *R*^2^ and its respective number of correlations yielded a value that more accurately represents the impact of observations towards *R*^2^ (narrower prediction bands). The number-weighted *R*^2^ metric indicates that *R*^2^ = 0 pairs have zero weight in predicting outcome. Correlations around *R*^2^ = 0.2 also revealed that most of the observational data is contributive towards lower *R*^2^ factors, and then a non-linear decrease closer to linearity is observed for higher *R*^2^. So lastly, Figure 15c also enriched our reductive attempts by pinpointing which weighted *R*^2^ lower threshold should be considered as meaningful. In the first iteration, a conservative *R*^2^ = 0.7 threshold was considered. This value was not arbitrary: it originated from the implementation of Pareto’s Principle as a lower threshold for a meaningful *R*^2^, in which 80% of the most significant correlations (*R*^2^ *>* 0.8) are expected to be expressed by 20% of all variables. This practical reasoning yielded a total of 236 correlations and an already drastic (96.3%) reduction in parameter complexity. By limiting the minimum number of literature reports required as *N* = 25 for an *R*^2^ = 0.7 to be significantly considered, the number of total correlations dropped to 33 correlations. However, most of these correlations pointed to variables of growth conditions relative to habitational markers, such as optimal temperature (*R*^2^ = 0.9) and specific yield (*R*^2^ = 0.8), which introduced correlational bias. The overreporting of habitational markers over structure and function attributes overshadows more desirable structural aspects (i.e. rheology) to be experimentally determined. Thus, in the second iteration we considered *R*^2^ = 0.8 but decreased parametrization to *N* = 5 as extreme-value conditions, outputting 58 correlations. From these, anionicity and structural archi-tecture were identified as consistent features in high *R*^2^ impact towards biological outcome. Heavy metal binding, gelling, viscosity, time-rheological properties and hydrodynamic radius were correlated with the percentage of uronic acids (*R*^2^ = 0.96), phosphate *R*^2^ = 0.89), sulfate (*R*^2^ = 0.83), as well as *α*(1→4) and *β*(1→4) linkages (*R*^2^ = 0.8–0.9). Although *N* is small, these compositional and structural behavior metrics are experimentally measurable. This parametric pre-processing was the basis of two publications which yielded corroborating experimental evidence [62, 63], hence validating the inherent value of a predictive analysis of `XPOL-DB` towards generating new insights.

**Figure 15:**
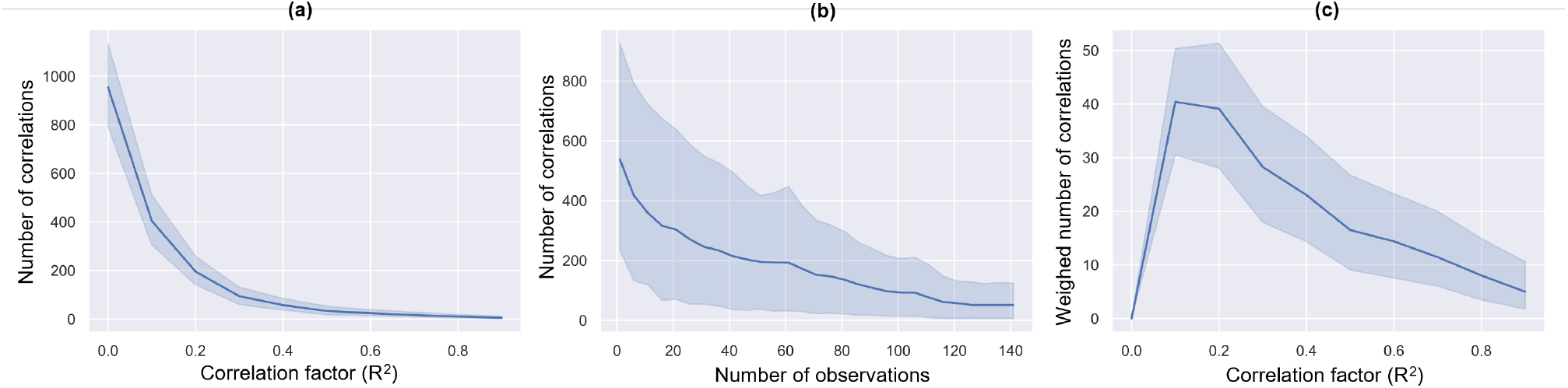
Summary of exploratory pair-plot correlation analysis. (**a**) Number of identified correlations as a function of *R*^2^. (**b**) Linear relationship between number of correlations as a function of number of recorded observations for each variable pair. (**c**) Weighed number of correlations as a function of *R*^2^ highlights bulk data reporting at *R*^2^ below 0.5.

**Table 4:** Cumulative *R*^2^ correlation distributions for variable pairs in `XPOL-DB`.

| Bins | Correlations | Cumulative (%) |
| --- | --- | --- |
| > 90% | 124 | 1.95 |
| > 80% | 171 | 2.69 |
| > 70% | 236 | 3.72 |
| > 60% | 310 | 4.88 |
| > 50% | 447 | 7.04 |
| > 40% | 656 | 10.33 |
| > 30% | 1019 | 16.04 |
| > 20% | 1692 | 26.64 |
| > 10% | 3070 | 48.33 |
| All | 6352 | 100.00 |

#### Monomeric Linearly-Additive Properties (MLAP)

Compared to monomer compositional data, the structural characteristics of extremophilic EPS is very sparsely reported. For instance, EPS net formal charge was reported 79.3% of the time, although most of these account for inductive reasoning from the ionic nature of the chromatographic column used during purification; followed by primary conformation (49.0%), linkage types (44.1%), secondary conformation (23.4%) and tertiary conformation (5.5%) reports. Therefore, to further infer on potential characteristics these EPS might express from a purely chemical standpoint but not standardly reported in the literature, an MLAP extrapolation methodology was developed, which generated new weighted-average metrics (Table 5). The MLAP calculations rely on a subset of *ab initio* assumptions:

**Table 5:**
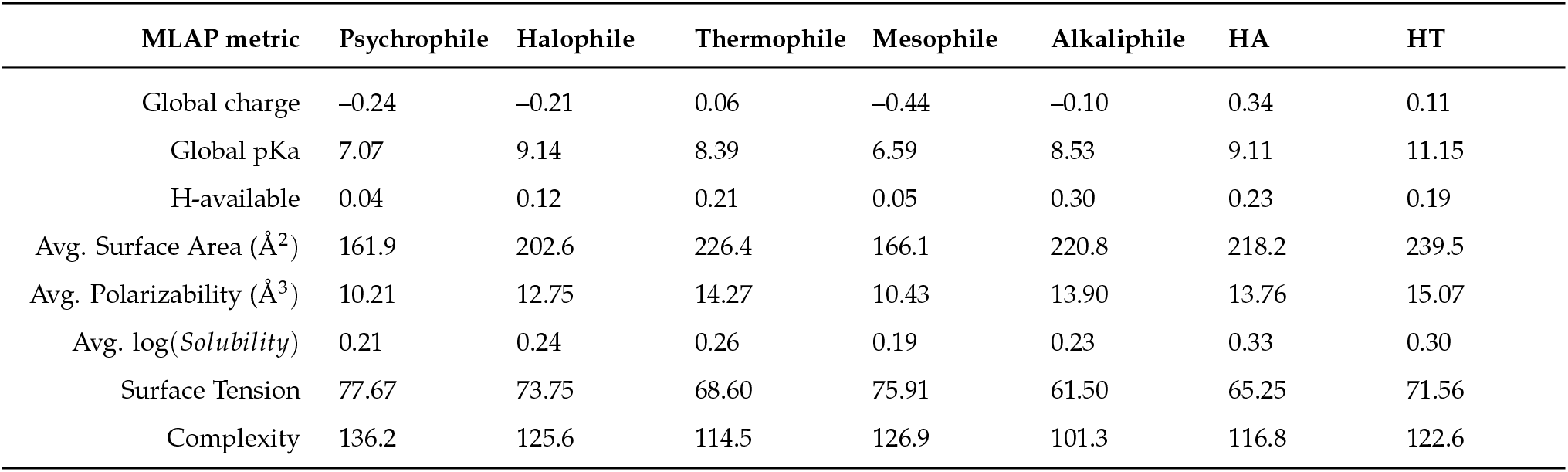
MLAP average metrics determined for each extremophilic type. The monomer data used for the calculations is based on theoretical predictions and experimental data cross-reported in *Pubchem, Chemspider*, and *Chemicalize* databases.

1. The chemical properties of each monomer in the extremophilic EPS structure are equal to their expressed properties in monomeric form.
2. The chemical properties of each monomer do not affect neighboring monomers.
3. Polymer traits are a linear combination of independent monomer traits.

The second assumption is overreaching, as it is known that, e.g. the type of established linkage hinges on the nature of the monomers participating in non-covalent interactions, and higher-order conformations arise from lower-order contributions from compositional parameters. Still, this simplification allowed, at the expense of some realism, to draw certain relativistic trends that corroborate real experimental data. Table 5 summarizes the MLAP properties that a given EPS SRU virtually possesses, such as linear approximations of global charge, global pKa, average topological polar surface area and polarizability, all these relevant in cryoprotective mechanisms as they reflect potential interactions at the polymer–ice interface. The extrapolatory validity of MLAP metrics was first confirmed by calculations of global charge, as EPS polarity was known from literature reports and could be cross-correlated. Indeed, psychrophilic (*−*0.24) and halophilic (*−*0.21) polysaccharides do demonstrate a negative global charge relative to the neutral thermophiles (0.06). Some mesophiles, such as FucoPol, had also shown cryoprotective traits [58, 59], hence the global charge observed (−0.44) also being consistent with observations in outcome. The global pKa of psychrophiles (7.1) is also lower than all other types (average 8.8), suggesting extensive deprotonation of their monomer residues, possibly arising from a higher incidence of uronic acids in psychrophilic structures. The average number of available H-atoms is a measure of hydrogen bond interactivity, with a lower value indicating greater hydrogen bond solvent interactions. Similar to global charge, psychrophiles (0.04) and halophiles (0.12) demonstrated higher solvent interactivity than other types, namely thermophiles (0.21). The average surface area (in Å^2^) and polarizability (in Å^3^) of polysaccharide chains, closely related to electrostatic traits and permanent dipole moments, showed lower values for psychrophiles (161.9 Å^2^, 10.21 Å^3^). A decreased solvent-accessible molecular surface area and volume is suggestive of conformational contraction. As an increased surface tension in psychrophiles (77.7 dyne/cm) and halophiles (73.8 dyne/cm) was observed relative to thermophiles (68.6 dyne/cm), this suggests that enhanced conformational packing with concomitant enhanced surface tension may be a requirement for kink-site adsorptive binding of cryoprotective molecules to crystal surfaces [114].

## Discussion

The first evidence of psychrophilic polysaccharides demonstrating ice-binding activity was the *Melosira arctica* algal EPS [15] and the *Colwellia psycherythraea* 34H bacterial CPS [26]. Initially, ice binding was ascribed to glycoproteins due to the absence of detailed structural information [115], but EPS production in microorganism survival strategies is of such signifi-cant recurrence [116] that deeper investigations were undertaken. Since then, accumulating knowledge on the known properties of extracellular polysaccharides (EPS) changed this paradigm: besides being the main constituents of biofilms and mediating nutrient uptake and cellular adhesion [117], their structural characteristics consistently showed traits of stress adaptability which pointed to their active participation in stress avoidance. Table 6 summarizes the current knowledge on structure–function relationships for extremophilic and mesophilic EPS trends, highlighting also the structural aspects of highest relevance to cryoprotective potential. In addition to the knowable universe presented in Table 6, the curation and multidimensional analysis of `XPOL-DB` revealed the following additional trends. Most polysaccharides have high molecular weight backbones up to 300 kDa, highly branched structures with an acidic nature imparted by uronic acids and show variable monomer compositions with rich diversity, usually composed of at least 2–5 different monomers, with psychrophilic polysaccharides having the highest SRU complexity amongst all extremophilic types. Depending on environmental pH, salinity and temperature, EPS can form complex architectural structures ranging from large networks of insoluble strands such as mucopolysaccharides often strengthened by covalent cross-linking and free Ca^2+^/Mg^2+^ ions, or hydrated gels that capitalize on functional group ionization to establish non-covalent interchain associations that confer strength and flexibility to the structure. The latter are often observed in sea ice and other subfreezing habitats. In an EPS structure, hydrogen bond interactions and electrostatic forces are the core predictors of bioactivity. Hydrophobicity often regulates cellular adhesion, allowing marine snow, microgels and biofilm structures to exist and thrive [118]. However, their intrinsic polyanionicity is a common factor in cryoprotection, heavy metal binding, salt tolerance and ROS scavenging mechanisms. This synergistic multifunctionality is often found in psychrophilic biofilms, whereas the constitutive polysaccharide architecture acts as a unitary defense mechanism against the cold.

**Table 6:**
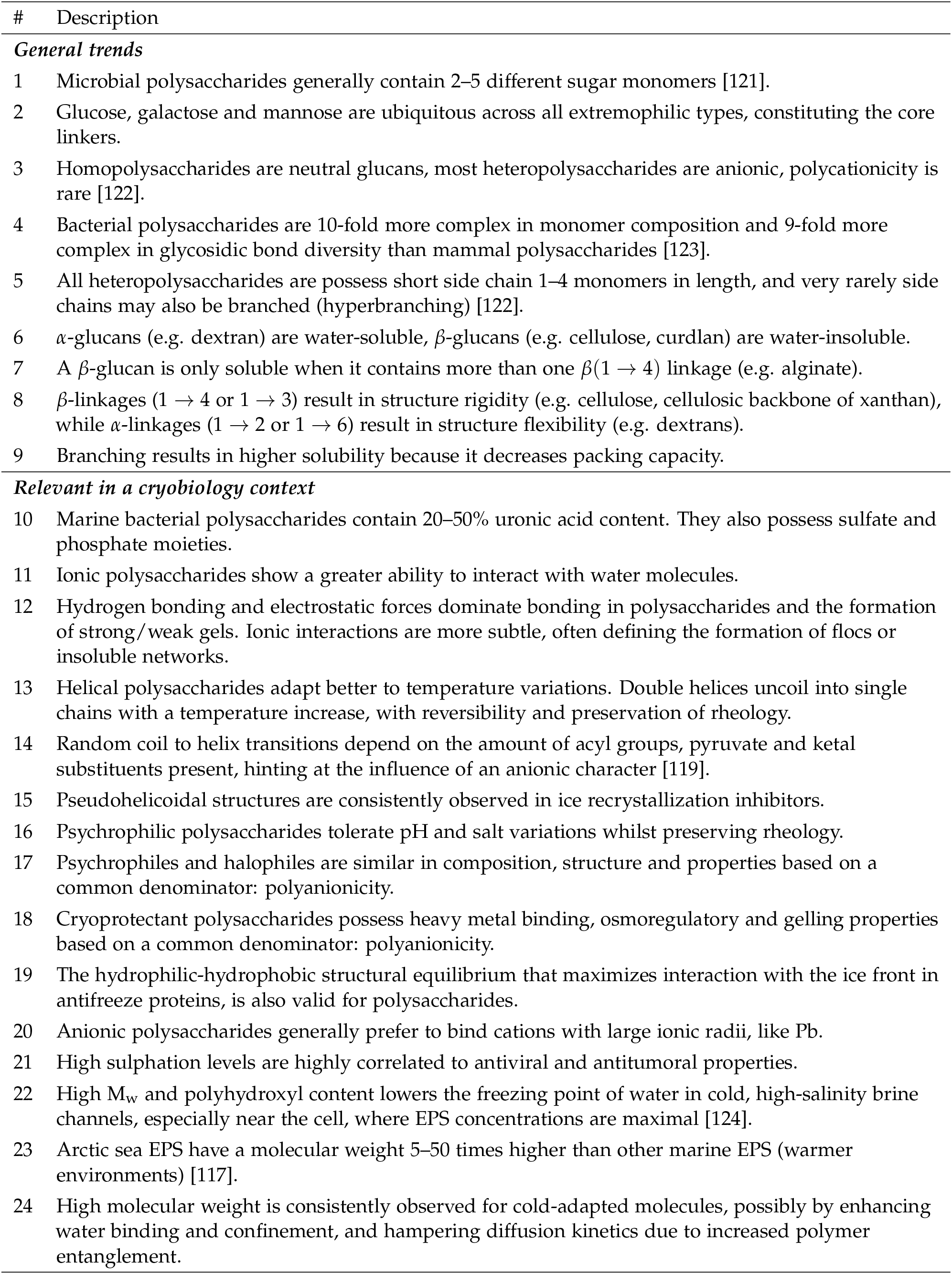
Summary of structure-function relationships found in extremophilic and mesophilic polysaccharides. The structural trends are divided in general, ubiquitously observed features for all polysaccharide structures, and psychrophile-specific observations of properties highly relevant for potential cryopreservation applications.

| # | Description |
| --- | --- |
| **General trends** |  |
| 1 | Microbial polysaccharides generally contain 2–5 different sugar monomers [121]. |
| 2 | Glucose, galactose and mannose are ubiquitous across all extremophilic types, constituting the core linkers. |
| 3 | Homopolysaccharides are neutral glucans, most heteropolysaccharides are anionic, polycationicity is rare [122]. |
| 4 | Bacterial polysaccharides are 10-fold more complex in monomer composition and 9-fold more complex in glycosidic bond diversity than mammal polysaccharides [123]. |
| 5 | All heteropolysaccharides are possess short side chain 1–4 monomers in length, and very rarely side chains may also be branched (hyperbranching) [122]. |
| 6 | α-glucans (e.g. dextran) are water-soluble, β-glucans (e.g. cellulose, curdlan) are water-insoluble. |
| 7 | A β-glucan is only soluble when it contains more than one $\beta(1 \rightarrow 4)$ linkage (e.g. alginate). |
| 8 | β-linkages ($1 \rightarrow 4$ or $1 \rightarrow 3$) result in structure rigidity (e.g. cellulose, cellulosic backbone of xanthan), while α-linkages ($1 \rightarrow 2$ or $1 \rightarrow 6$) result in structure flexibility (e.g. dextrans). |
| 9 | Branching results in higher solubility because it decreases packing capacity. |
| **Relevant in a cryobiology context** |  |
| 10 | Marine bacterial polysaccharides contain 20–50% uronic acid content. They also possess sulfate and phosphate moieties. |
| 11 | Ionic polysaccharides show a greater ability to interact with water molecules. |
| 12 | Hydrogen bonding and electrostatic forces dominate bonding in polysaccharides and the formation of strong/weak gels. Ionic interactions are more subtle, often defining the formation of flocs or insoluble networks. |
| 13 | Helical polysaccharides adapt better to temperature variations. Double helices uncoil into single chains with a temperature increase, with reversibility and preservation of rheology. |
| 14 | Random coil to helix transitions depend on the amount of acyl groups, pyruvate and ketal substituents present, hinting at the influence of an anionic character [119]. |
| 15 | Pseudohelical structures are consistently observed in ice recrystallization inhibitors. |
| 16 | Psychrophilic polysaccharides tolerate pH and salt variations whilst preserving rheology. |
| 17 | Psychrophiles and halophiles are similar in composition, structure and properties based on a common denominator: polyanionicity. |
| 18 | Cryoprotectant polysaccharides possess heavy metal binding, osmoregulatory and gelling properties based on a common denominator: polyanionicity. |
| 19 | The hydrophilic-hydrophobic structural equilibrium that maximizes interaction with the ice front in antifreeze proteins, is also valid for polysaccharides. |
| 20 | Anionic polysaccharides generally prefer to bind cations with large ionic radii, like Pb. |
| 21 | High sulphation levels are highly correlated to antiviral and antitumoral properties. |
| 22 | High $M_w$ and polyhydroxyl content lowers the freezing point of water in cold, high-salinity brine channels, especially near the cell, where EPS concentrations are maximal [124]. |
| 23 | Arctic sea EPS have a molecular weight 5–50 times higher than other marine EPS (warmer environments) [117]. |
| 24 | High molecular weight is consistently observed for cold-adapted molecules, possibly by enhancing water binding and confinement, and hampering diffusion kinetics due to increased polymer entanglement. |

### Contributions to extremophilic structure-function relationships

A psychrophilic microenvironment may confer protection against freezing damage in glacial water habitats or desiccation in glacial ice, buffer against highly saline environments found in permafrost brine, quench free radicals and chelate heavy metals in solution while also scavenging biologically beneficial nutrients, and aid in cell motility in adherence in the host biofilm [119]. The very high similarity be-tween psychrophilic and halophilic polysaccharides arises from a shared stress adaptation. Ice formation involves solute rejection and supersaturation, creating a brine-like medium commonly found in Arctic ice channels and pore spaces. In these conditions, psychrophiles experience halophilic survival conditions, to which they also must adapt to in order to survive [120]. While at a macroscale this dual stressor creates the necessity for hybrid adaptation, at a microscale both ice growth and high salinity share the same fundamental mechanisms: diffusion kinetics and membrane regulation. While it remains valid how natural selection and an interconnected ecosystem of different biomes (sea ice, brine channels, sea water, etc.) promoted chemical adaptation towards survival, it is intriguing how the mesophile *Enterobacter* A47 produces a fucose-rich polysaccharide (FucoPol) with proven ice growth inhibition and a versatile structure capable of tolerating increased salinity [58, 59]. Nevertheless, its multifunctionality reinforces the proposition that certain polysaccharides can express a dualistic defense mechanism due to compositional and structural features alone, notwithstanding that environmental origin predominantly drives such phenotypical adaptation. Out of more than 6000 correlations, two main parametric denominators proved the most impactful in potentially predicting a cryoprotective outcome: structural polyanionicity and an influence in molecular diffusion. High likelihood predictors of cryoprotective potential are monomer compositions skewed towards elevated uronic acid content, conformations resembling helical structures, increased molecular weight, rheological properties inducive of high viscosity and gelling behavior and secondary properties that rely on anionic interactions to exert an effect, such as antioxidant potential or heavy metal binding.

### Concluding remarks

Here, a cryopreservation-centric database of extremophilic polysaccharide structures, `XPOL-DB`, was compiled. The aggregated view of extremophilic polysaccharide research, clustering of structure-function relationships by extremophilic type as a measure of relating their defensive chemical adaptation to unique external stressors, and superposition of psychrophilic and thermophilic profiles, allowed us to differentially highlight the impact of divergent evolutionary biology needs in phenotypical polysaccharide chemistry. This systematic analysis in particular focused on the cryoprotective effect. We elucidated the highest likelihood structure-function relationships that are expressed by extremophilic EPS towards stress-response, particularly freezing injury.

From these insights, we set out to define from a practical standpoint a subset of experimental variables that could quickly inform the cryobiologist if a given polysaccharide has the potential for cryoprotection. The validity of this analytical effort was later corroborated in parallel studies that leveraged the novel insights gathered herein. A systematic review of this sort allowed to extend knowledge on all known structure-function relationships, either by recalling and validating previous literature trends, or by generating new insights from hidden correlations in data. A pre-experimental *in silico* investigation can accelerate laboratory design-of-experiment by statistically screening for the most impactful, measurable variables towards a cryoprotective outcome, thereby also potentiating future reverse engineering research of relevant polysaccharide structures through predictive analysis of their functionality.

## CRediT Authorship

B.M.G. conceptualized study, performed experiments, data analysis, wrote manuscript. F.F., J.L., J.S. reviewed manuscript, supervised, funded. All authors have read and agreed to the published version of the manuscript.

## Data Availability Statement

The data that support the findings of this study are available from the corresponding author upon request.

