## Supplementary Information for "Polysaccharides in cryopreservation: multidimensional systematic review of extremophilic traits and the role of selective pressure in structure-function relationships"

### **1. XPOL-DB database documentation**

Table SI.1 shows the full list of extremophilic strains included in XPOL-DB. The full database can be found [online here](https://docs.google.com/spreadsheets/d/1Pau7IhcXFqHlOs0PBtbPr0CBMyV3wPbn/edit?usp=sharing&ouid=110571406079464345723&rtpof=true&sd=true), with respective references to literature. We often encountered knowledge gaps in during the systematic analysis. Biometrics data was collected from the original source paper, but in cases that data was not presented, it was retrieved from secondary sources to ensure statistical continuity, by the following order of priority:

1. Original source paper;
2. Search for strain characterization in *International Journal of Systematic and Evolutionary Microbiology,* which often contained information on taxonomy, phenotype, growth conditions and heavy metal tolerance.
3. Search for strain in depositories, such as *Bergey’s Manual of Systematic Bacteriology*.

Table SI.1 – Full list of extremophilic strains and their type, included in XPOL-DB.

| ID | Strain | ExtremeType | ID | Strain | ExtremeType |
| --- | --- | --- | --- | --- | --- |
| 1 | *Geobacillus thermodenitrificans* ArzA-6 [1] | Thermophile | 74 | *Halomonas almeriensis* M8^T^ [2] | Halophile |
| 2 | *Geobacillus toebii* ArzA-8 [1] | Thermophile | 75 | *Halomonas almeriensis* M8^T^ [2] | Halophile |
| 3 | *Rhodothermus marinus* DSM4252^T^ [3] | Thermophile | 76 | *Vibrio sp.* QY101 [4] | Halophile |
| 4 | *Rhodothermus marinus* DSM4252^T^ [3] | Thermophile | 77 | *Halomonas stenophila* B100 [5] | Halophile |
| 5 | *Rhodothermus marinus* DSM4252^T^ [3] | Thermophile | 78 | *Halomonas stenophila* N12^T^ [5] | Halophile |
| 6 | *Rhodothermus marinus* DSM4252^T^ [3] | Thermophile | 79 | *Salipiger mucosus* A3^T^ [6] | Halophile |
| 7 | *Rhodothermus marinus* MAT493 [3] | Thermophile | 80 | *Idiomarina fontislapidosi* F23^T^ [7] | Halophile |
| 8 | *Rhodothermus marinus* MAT493 [3] | Thermophile | 81 | *Idiomarina fontislapidosi* F23^T^ [7] | Halophile |
| 9 | *Rhodothermus marinus* MAT493 [3] | Thermophile | 82 | *Idiomarina ramblicola* R22^T^ [7] | Halophile |
| 10 | *Geobacillus sp.* TS3-9 [8] | Thermophile | 83 | *Idiomarina ramblicola* R22^T^ [7] | Halophile |
| 11 | *Aeribacillus pallidus* 418 [9–11] | Thermophile | 84 | *Alteromonas hispanica* F32^T^ [7] | Halophile |
| 12 | *Aeribacillus pallidus* 418 [9–11] | Thermophile | 85 | *Halomonas eurihalina* F2-7 [12,13] | Halophile |
| 13 | *Brevibacillus thermoruber* 423 [14] | Thermophile | 86 | *Halomonas ventosae* A112^T^ [15] | Halophile |
| 14 | *Anoxybacillus sp.* R4-33 [16] | Thermophile | 87 | *Halomonas ventosae* A116 [15] | Halophile |
| 15 | *Aeribacillus pallidus* YM-1 [17] | Thermophile | 88 | *Halomonas anticariensis* FP35^T^ [15] | Halophile |
| 16 | *Thermus aquaticus* YT-1 [18] | Thermophile | 89 | *Halomonas anticariensis* FP36 [15] | Halophile |
| 17 | *Geobacillus thermodenitrificans* B3-72 [19,20] | Thermophile | 90 | *Halomonas maura* S-30 [21] | Halophile |
| 18 | *Geobacillus thermodenitrificans* B3-72 [19,20] | Thermophile | 91 | *Aphanothece halophytica* GR02 [22] | Halophile |
| 19 | *Geobacillus tepidamans* V264 [23] | Thermophile | 92 | *Aphanothece halophytica* GR02 [22] | Halophile |
| 20 | *Geobacillus sp.* 4004 [24] | Thermophile | 93 | *Aphanothece halophytica* GR02 [22] | Halophile |
| 21 | *Geobacillus sp.* 4004 [24] | Thermophile | 94 | *Aphanothece halophytica* GR02 [22] | Halophile |
| 22 | *Geobacillus sp.* 4004 [24] | Thermophile | 95 | *Aphanothece halophytica* GR02 [22] | Halophile |
| 23 | *Bacillus thermoantarcticus* EPS1 [25] | Thermophile | 96 | *Aphanothece halophytica* GR02 [22] | Halophile |
| 24 | *Bacillus thermoantarcticus* EPS2 [25] | Thermophile | 97 | *Halomonas alkaliantarctica* CRSS [26] | Haloalkaliphile |
| 25 | *Bacillus licheniformis* B3-15 [27–29] | Halothermophile | 98 | *Halomonas alkaliantarctica* CRSS [26] | Haloalkaliphile |
| 26 | *Pseudoalteromonas sp.* MER144 [30] | Psychrophile | 99 | *Bacillus spp.* [31,32] | Haloalkaliphile |
| 27 | *Lactobacillus sakei* TMW 1.411 [33] | Psychrophile | 100 | *Cronobacter sakazakii* [34] | Alkaliphile |
| 28 | *Lactobacillus sakei* TMW 1.411 [33] | Psychrophile | 101 | *Bacillus cereus* [35] | Alkaliphile |
| 29 | *Lactobacillus sakei* TMW 1.411 [33] | Psychrophile | 102 | *Bacillus thuringiensis* [35] | Alkaliphile |
| 30 | *Winogradskyella sp.* CAL384 [36] | Psychrophile | 103 | *Proteus mirabilis* | Alkaliphile |
| 31 | *Winogradskyella sp.* CAL396 [36] | Psychrophile | 104 | *Enterobacter sp.* A47 FucoPol | Mesophile |
| 32 | *Colwellia sp.* GW185 [36] | Psychrophile | 105 | *Marinobacter strain* W1–16 [42] | Psychrophile |
| 33 | *Shewanella sp.* CAL606 [36] | Psychrophile | 106 | *Pseudomonas strain* UC-1 [43] | Psychrophile |
| 34 | *Colwellia psychrerythraea* 34H-Ala [44,45] | Psychrophile | 107 | *Pseudoalteromonas strain* S-15-13 EPS-II [46–48] | Psychrophile |
| 35 | *Colwellia psychrerythraea* 34H-Thr [49] | Psychrophile | 108 | *Pseudoalteromonas arctica* KOPRI 21653 [50] | Psychrophile |
| 36 | *Colwellia psychrerythraea* 34H-CPS [51] | Psychrophile | 109 | *Psychrobacter arcticus* 273-4 LPS [52,53] | Psychrophile |
| 37 | *Pseudoalteromonas elyakovii* Arcpo 15 [54] | Psychrophile | 110 | *Psychrobacter arcticus* 273-4 Mannan [55] | Psychrophile |
| 38 | *Pseudomonas sp.* ID1 [56–58] | Psychrophile | 111 | *Bacillus thuringiensis* YY529 [59] | Mesophile |
| 39 | *Cobetia marina* DSMZ 4741 [60] | Halophile | 112 | *Upis ceramboides* Xylomannan [61] | Psychrophile |
| 40 | *Polaribacter sp.* SM1127 [62] | Psychrophile | 113 | *Flammulina velutipes* Xylomannan (mycel.) [63,64] | Psychrophile |
| 41 | *Pseudoalteromonas sp.* SM20310 [65] | Psychrophile | 114 | *Flammulina velutipes* Xylomannan (body) [63,64] | Psychrophile |
| 42 | *Pseudoalteromonas sp.* S-5 [66] | Psychrophile | 115 | *Lycium barbarum* EPS [67–69] | Mesophile |
| 43 | *Pseudoalteromonas sp.* SM9913 [70] | Psychrophile | 116 | *Astragalus membranaceus* EPS [71,72] | Mesophile |
| 44 | *Flavobacterium frigidarium* CAM005 [73] | Psychrophile | 117 | *Acinetobacter lwoffii* EK30A [74] | Psychrophile |
| 45 | *Myroides odoratus* CAM030 [73] | Psychrophile | 118 | *Acinetobacter sp.* VS-15 [75,76] | Psychrophile |
| 46 | *Polaribacter irgensii* CAM006 [73] | Psychrophile | 119 | *Acinetobacter lwoffii* EK67 [75,76] | Psychrophile |
| 47 | *Pseudoalteromonas sp.* CAM003 [73] | Psychrophile | 120 | *Psychrobacter cryohalolentis* K5^T^ [77,78] | Psychrophile |
| 48 | *Pseudoalteromonas sp.* CAM015 [73] | Psychrophile | 121 | *Psychrobacter muricolla* 2pS^T^ [79] | Psychrophile |
| 49 | *Pseudoalteromonas sp.* CAM023 [73] | Psychrophile | 122 | *Moritella viscosa* M2-226 [80,81] | Psychrophile |
| 50 | *Pseudoalteromonas sp.* CAM025 [73,82,83] | Psychrophile | 123 | *Flexibacter psychrophilum* 259-93 [84,85] | Psychrophile |
| 51 | *Pseudoalteromonas sp.* CAM036 [73] | Psychrophile | 124 | *Idiomarina zobellii* KMM 231^T^ [86,87] | Halophile |
| 52 | *Pseudoalteromonas sp.* CAM064 [73] | Psychrophile | 125 | *Psychromonas arctica* LPS [88,89] | Psychrophile |
| 53 | *Shewanella livingstonensis* CAM090 [73] | Psychrophile | 126 | *Pseudomonas sp.* BGI-2 [90] | Psychrophile |
| 54 | *Pseudoalteromonas haloplanktis* TAC 125 [91–93] | Psychrophile | 127 | *Zygosaccharomyces rouxii* EPS-3791 [94] | Halophile |
| 55 | *Pseudoalteromonas haloplanktis* TAB 23 [91,92] | Psychrophile | 128 | *Laminaria japonica* LJP-P3 [95–97] | Mesophile |
| 56 | *Pseudomonas sp.* NCMB 2021 [98] | Psychrophile | 129 | *Rhodiola rosea* EPS [99–101] | Mesophile |
| 57 | *Pseudomonas sp.* NCMB 2021 [98] | Psychrophile | 130 | *Gynostemma Pentaphyllum* GPP1-α [102–106] | Mesophile |
| 58 | *Halomonas nitroreducens* WB1 [107] | Halothermophile | 131 | *Lemna minor* Lemnan [108–112] | Psychrophile |
| 59 | *Halomonas nitroreducens* WB1 [107] | Halothermophile | 132 | *Comarum palustre* Comaruman [108,113] | Psychrophile |
| 60 | *Halomonas nitroreducens* WB1 [107] | Halothermophile | 133 | *Bergenia classifolia* Bergenan [108,114,115] | Psychrophile |
| 61 | *Bacillus licheniformis* T14 [116–118] | Halothermophile | 134 | *Potamogeton natans* Potamogetonan [108] | Psychrophile |
| 62 | *Geobacillus sp.* 1A60 [119] | Halothermophile | 135 | *Tanacetum vulgare* Tanacetan [108,120,121] | Psychrophile |
| 63 | *Chromohalobacter canadensis* 28 [122] | Halophile | 136 | *Rauwolfia serpentina* Rauwolfian [121,123,124] | Psychrophile |
| 64 | *Halolactibacillus miurensis* SEEN MKU3 [125] | Halophile | 137 | *Heracleum sosnowskyi* Heracleum [121,126] | Psychrophile |
| 65 | *Kocuria rosea* ZJUQH [127] | Halophile | 138 | *Scotiellopsis terrestris* St [108] | Psychrophile |
| 66 | *Vibrio alginolyticus* CNCM I-4994 [128] | Halophile | 139 | *Nostoc muscorum* Nm [108,129] | Psychrophile |
| 67 | *Halomonas smyrnensis* AAD6^T^ [130–133] | Halophile | 140 | AU-701 Apple pectin [134] | Mesophile |
| 68 | *Alteromonas macleodii* [135] | Mesophile | 141 | *Salvia miltiorrhiza* EPS [136–140] | Psychrophile |
| 69 | *Alteromonas infernus* | Mesophile | 142 | *Aloe arborescens* AA3 Pectin [141] | Mesophile |
| 70 | *Alteromonas sp.* 1644 [142] | Mesophile | 143 | *Hericium erinaceus* BP 16 [143–145] | Mesophile |
| 71 | *Alteromonas sp.* 1545 | Mesophile | 144 | *Phoma herbarum* CCFEE 5080 EPS [146] | Psychrophile |
| 72 | *Pseudoalteromonas sp.* 721 | Mesophile | 145 | *Bacillus enclensis* AP-4 [147,148] | Halophile |
| 73 | *Vibrio diabolicus CNCM I-1629* [151-152] | Mesophile |  |  |  |

### **2. A primer on polysaccharides**

A carbohydrate is a molecule composed of carbon, hydrogen and oxygen that follows the empirical formula C*_x_*(H_2_O)*_y_*, where *x* and *y* ≥ 3. Because hydrogen and oxygen are present in the same proportions as in water, carbohydrates have been described as hydrated forms of carbon [149]. Currently, the term carbohydrate is more commonly used to refer to linear or cyclical structures of five or six-carbon monosaccharides, which can polymerize by condensation reactions and form oligo (3–10 monomers) or polysaccharides (10+ monomers), linked by glycosidic linkages. In their cyclical form, the hydroxyl groups linked to each carbon can be positioned above or below the axial plane (Figure SI.1).

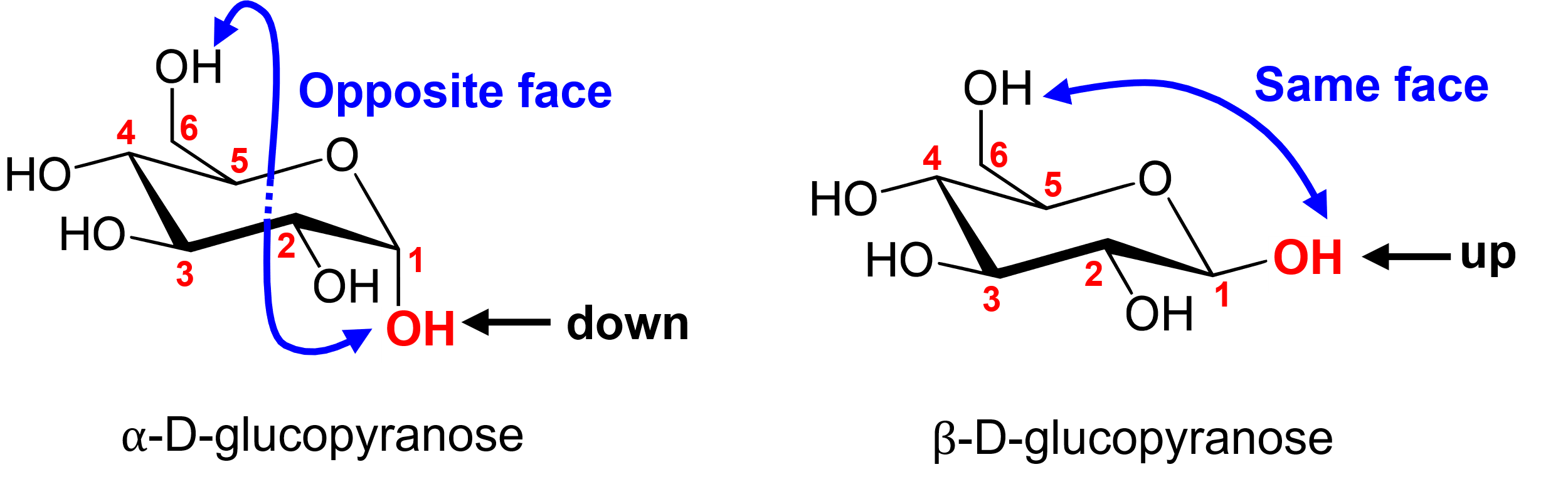

Figure SI.1 – Monosaccharide linkage definition relative to the anomeric carbon of D-glucose.

These combinations are what define the many different sugar monomers known, *i.e.* glucose (Glc), galactose (Gal), mannose (Man), among others. The anomeric carbon C_1_ participates in bonding and defines the type of glycosidic linkage. If the –OH group at C_1_ is in an axial position, an α-linkage emerges. Otherwise, the equatorial position defines a β-linkage. The definition of a carbohydrate polymer spans many types of sugar-based structures. This paper focuses on polysaccharides, specifically those of biological origin, which can be extracellularly secreted polysaccharides (EPS) or capsular polysaccharides (CPS). Bacterial EPS are often produced inside the microorganism via the Wzx/Wzy or Synthase-dependent pathways and then excreted into the medium to form a biofilm, although some EPS (*e.g.* dextran, levan) are directly synthesized outside the cell by glycosyltransferases covalently linked to the cell surface via the GT-dependent pathway [150]. A CPS is produced in a bound state to the outer bacterial wall via the ABC transporter-dependent pathway, especially in *Gram*-negative bacteria, and is usually of pathogenic nature, playing a role in the immune response [150]. In the context of cryobiology, AFGPs are common examples of glycosylated polymers.
